# Machine-Guided Dual-Objective Protein Engineering for Deimmunization and Therapeutic Functions

**DOI:** 10.1101/2025.02.24.639918

**Authors:** Eric Wolfsberg, Jean-Sebastien Paul, Josh Tycko, Binbin Chen, Michael C. Bassik, Lacramioara Bintu, Ash A. Alizadeh, Xiaojing J. Gao

## Abstract

Cell and gene therapies often rely on the expression of exogenous proteins derived from nonhuman organisms. An emerging consensus is to reduce the potential immunogenicity of such therapies by instead using human protein domains. However, as we engineer these human-derived proteins, we create nonhuman peptides at the linkers or junctions between domains and at mutated residues within them, which still pose a risk of immunogenicity that has largely been left unaddressed. Here, we present a modular workflow to simultaneously optimize the functions of proteins and minimize their immunogenic risk using existing machine learning models that predict protein function and nonhuman peptide immunogenicity from their sequences. We first applied this workflow to existing transcriptional activation and bio-orthogonal RNA binding domains. Then we generated a set of small molecule-controllable transcription factors with human-derived zinc finger DNA-binding domains for targeting orthogonal non-genomic DNA sequences. Finally, we established a workflow for creating deimmunized zinc finger arrays to target arbitrary genomic DNA sequences and used it to upregulate expression of two therapeutically relevant genes, UTRN and SCN1A. Our future-proof, modular workflow offers a proof of principle for making cell and gene therapies safer and more efficacious through dual-objective protein optimization using state-of-the-art algorithms.

## Introduction

Cell and gene therapies are an emerging pillar of modern medicine. However, as it is routine to use nonhuman proteins in these contexts, either derived from other organisms or engineered from native human proteins, they have the potential to be immunogenic. This has been seen in therapies involving both chimeric antigen receptor (CAR)-expressing T cells^1–3^ and gene editors using CRISPR-associated proteins.^4–8^ When peptides from these nonhuman proteins are presented on cell surfaces by major histocompatibility complex (MHC) molecules, the peptide-MHC complexes may activate receptors on T cells and induce adaptive immune responses. While there are efforts to remove such MHC epitopes *post hoc*, there has been a lack of concerted effort to “humanize” proteins used in cell and gene therapies. This is in contrast to the shift in therapeutic monoclonal antibodies (mAbs) from murine to chimeric to fully human with a corresponding decrease in immunogenicity.^9–11^ Human proteins, in principle, don’t share the risk of adaptive immune responses because negative selection in T cell development should eliminate T-cell receptors (TCRs) specific to self-antigens from human proteins.^12^ This suggests the strategy of replacing nonhuman proteins in these contexts with ones that exclusively use domains either sourced from or similar to ones in the human proteome. Recent examples of this strategy have emerged in the design of RNases,^13^ transcription factors,^14^ synthetic receptors,^15^ and proteases.^16^

However, the immunogenicity risk is not entirely eliminated by the exclusive use of human protein domains (**Fig. 1a**). When an engineered protein fuses domains which do not naturally occur next to one another, the junctions between these domains will be degraded into novel peptides which can, in principle, be loaded onto MHC molecules and be bound by T-cell receptors. A similar argument can be made for single-domain proteins with point mutations that modify their activity for purposes such as bio-orthogonalization: peptides containing mutated amino acid residues are not fully human, and thus can be bound by TCRs which are not selected against. Evidence exists that analogous natural phenomena, such as alternative splicing^17–20^ and tumor-associated point mutations^21^ can cause immune responses to human-derived proteins. Additionally, the primary contributing factor to the immunogenicity of fully human mAbs has been suggested to be the complementarity-determining sequence,^22^ indicating that even short non-self sequences can suffice to induce an immune response against therapeutic proteins.

**Fig. 1.**
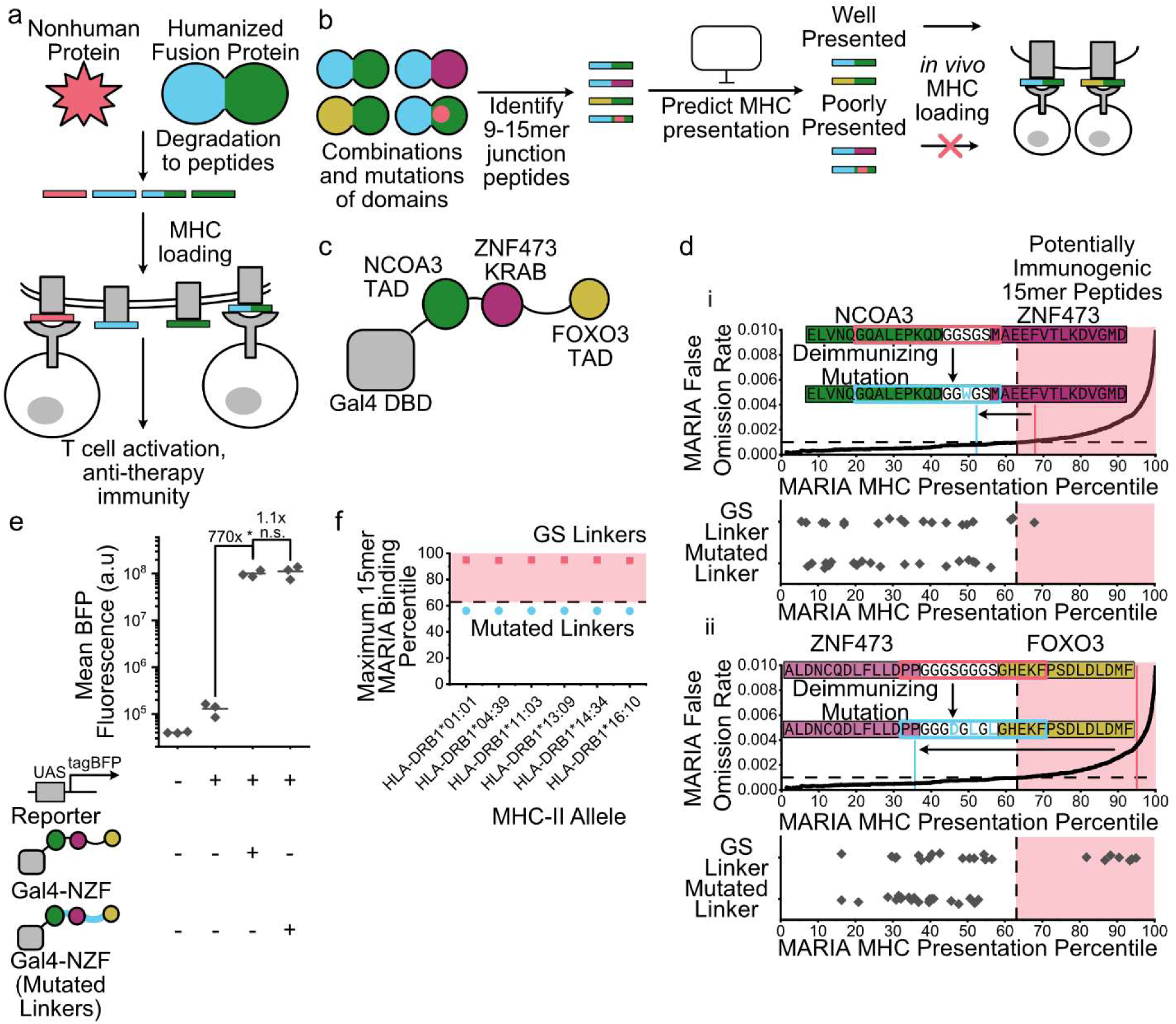
Deimmunization workflow and application to NZF transcriptional activation domain. (**a**) Schematic of adaptive anti-therapeutic immunity against therapeutic nonhuman proteins and human-derived fusion proteins. (**b**) Schematic of deimmunization workflow. (**c**) Schematic of NZF transcriptional activation domain attached to Gal4 DNA-binding domain. (**d**) Deimmunization of NZF’s NCOA3-ZNF473 (i) and ZNF473-FOXO3 (ii) linkers through the introduction of mutations which reduce the MARIA percentile score of all peptides within those linkers to below the threshold corresponding to a false omission rate of 0.001. 15mer peptide identities and MARIA percentile scores are in **Supp. Table S5**. (**e**) Flow cytometric comparison of NZF transcriptional activation with GS and mutated linkers. Unless otherwise stated, flow cytometry data occurs following transient transfection into HEK293 cells, data points represent the geometric means of cells gated for high transfection for each of three biological replicates (see Methods), lines represent arithmetic means of each set of biological replicates, fold changes represent the ratios of those means across conditions, and significance is tested by a Bonferroni-corrected two-tailed Welch’s *t-*test. * denotes p < 0.05 and n.s. denotes p > 0.05. (**f**) Maximum MARIA scores for GS and mutated linker sequences across a selection of MHC-II alleles. Data for all HLA-DR alleles which can be estimated by MARIA are in **Supp. Table S2**.

While the use of human domains in therapeutic proteins is only just emerging, the potential immunogenicity derived from junctions and mutations is particularly less appreciated in cell and gene therapies. This may be due to the relative novelty of these therapies, but the adoption of engineering practices that reduce potential immunogenicity in the early stages of development can both save costs and reduce patient risk compared to considering it only after a complete therapy is developed. Therefore, we propose and demonstrate in this paper a framework for eliminating potentially immunogenic peptide sequences from engineered human-derived proteins. This is a complex task involving both the prediction of MHC binding and the optimization of protein function, both fast-moving fields. Because of this, we choose to emphasize the modularity of our workflow rather than the outcomes alone. While we need to use particular existing algorithms to demonstrate the workflow, these algorithms can be readily swapped out for alternatives, including superior ones which we have no doubt will materialize in the future. In this way we can bring together work in other fields to empower the creation of efficacious and safe cell and gene therapeutics.

We chose the engineering of DNA-binding domains and transcription factors (TFs) as an especially pertinent and challenging test case. Broadly speaking, there are two classes of therapeutically relevant DNA binding domains and corresponding binding sites. The first class is those which bind to a motif absent from the human genome to regulate the transcription of a transgene in a controllable manner. These include both bacterially-derived domains such as TetR and derivatives thereof such as rtTA^23^ and more human-like domains such as zinc finger (ZF) arrays that are targeted to a particular non-genomic DNA motif by mutagenesis and directed evolution.^14,24^ The second class is those which bind selectively to a specific motif within the human genome so as to affect that site in some way, either by cleaving it or affecting transcription. These include most obviously CRISPR-associated proteins which are targeted using guide RNAs^25^, as well as other domains such as TALEs^26^ or ZFs^27,28^ which are designed or evolved to bind to genomic sites. For both these cases, novel DNA-binding activity has to come from nonhuman proteins, either in the form of domains from other organisms such as TetR and Cas9 or human-like domains such as ZFs which must be mutated to exhibit binding specificity distinct from existing human proteins. The use of nonhuman DNA-binding domains has empirically been implicated in undesirable immunogenicity,^7,29–31^ while mutated ZF arrays likewise have the potential for immunogenicity due to the presence of nonhuman peptides created during the evolution process. In the bulk of this paper, we examined whether we could program DNA-binding domains to meet the dual objectives of functionality and deimmunization, combining several state-of-the-art machine learning models.

## Results

We envision a workflow for generating novel, deimmunized, human-derived protein domains (**Fig. 1b**). First, we collect candidate protein sequences based on their predicted functionality. For fusion proteins, this collection may consist of alternative sets of protein domains to be fused together (for example, the blue-green, yellow-green, and blue-purple pairs in **Fig. 1b**). It may also include mutations applied to one or more fixed protein domains, introduced either to modify the function of the protein or to deimmunize it (for example, the blue-green pair with a red mutation in **Fig. 1b**). From each of these candidates a set of nonhuman peptides is extracted, consisting of peptides that cross the junction between human domains, including nonhuman interdomain linkers, and peptides that contain artificial mutations. The ability of these peptides to be presented by one or more MHC alleles is then computationally predicted. If a particular candidate has no nonhuman peptides that are predicted to be presented by the MHC alleles, conditional on the accuracy of the prediction algorithm, it could dramatically reduce the likelihood of an anti-therapeutic immune response.

A key aspect of this workflow is that it leaves open the choice of algorithm to predict MHC presentation, the choice of threshold to distinguish presented from non-presented peptides, and the choice of which MHC classes and alleles to analyze. Many algorithms exist for the prediction of peptide presentation by both Class I^32–40^ and Class II^34,35,41–44^ MHC molecules, and those seeking to adapt this workflow for their own purposes may freely slot in whichever they judge to be the best. It is worth noting that many use such algorithms to identify highly-presented peptides with the goal of using such peptides in immunotherapies that provoke an immune response.^45^ Due to that limitation of existing models we recommend relatively strict thresholding to exclude peptides that are potentially presented by the MHC. The examples shown below all conform to a threshold corresponding to a false omission rate (FOR) of 0.001, that is, at which if 1000 peptides are predicted not to be presented only 1 of those will, on average, in fact be presented by the MHC. This threshold is beyond adequate because few, if any, proteins created using this approach will contain close to 1000 nonhuman peptides. Finally, while in principle this workflow is applicable for prediction of both MHC Class I and Class II presentation, here we choose to focus on MHC Class II presentation. This is because although the major risk of anti-therapy immunity against engineered cells is cytotoxicity from MHC-I-binding CD8+ T cells, assistance from CD4+ T cells is also necessary for activating cytotoxic T cells.^46,47^ Moreover, there is evidence that MHC-II presentation is more strongly selected against in tumors than MHC-I presentation.^48^ This implies that the survival of tumor cells, and presumably that of engineered cells as well, relies more on the avoidance of MHC-II than MHC-I antigen presentation. We chose to predict MHC presentation primarily using MARIA,^41^ a relatively recent MHC-II prediction algorithm trained on mass spectrometry-derived antigen data as well as conventional binding measurements. Most of our analysis focuses on the MHC-II allele HLA-DRB1*01:01, one of the 10 most common DRB1 alleles across global populations,^49^ though we have observed a relatively weak effect of MHC-II allele on the prediction of peptide presentation (**Fig. 1f**, **Fig. 6h**).

As a simple example of our deimmunizing workflow, we show its application to the potent NZF transcriptional activation domain, which consists of activation domains from the human proteins NCOA3, ZNF473, and FOXO3.^50^ While NZF, in contrast to the previously commonly used VPR domain,^51^ has the benefit of consisting entirely of human domains, these domains are linked by nonhuman glycine-serine (GS) linkers (**Fig. 1c**). These linkers, when combined with the adjoining portions of the activation domains, contain 15mer peptides that are predicted by MARIA to be presented by HLA-DRB1*01:01 (**Fig. 1d**) and other MHC-II alleles (**Fig. 1f**). To eliminate these peptides, we exhaustively generated variants of these linkers with increasing numbers of mutations. We found that for the linker between the NCOA3 and ZNF473 domains one mutation (S3W) reduced the predicted presentation of all nonhuman peptides to below the false omission rate threshold, which for MARIA lies at about the 63^rd^ percentile of peptide scores (**Fig. 1di**). Three mutations (S4D, G6L, and S8L) were necessary to do the same for the linker between ZNF473 and FOXO3 domains (**Fig. 1dii**). When fused to the Gal4 DNA-binding domain, this deimmunized version of the NZF domain could induce transcription of BFP from a co-transfected reporter plasmid to a level equal to the original version (**Fig. 1e**). This means that deimmunization of the linkers within NZF did not impact its function for transcriptional activation. Moreover, this combination of mutations reduced the MHC-presentation score of all linker peptides below the desired threshold not only for the HLA-DRB1*01:01 allele for which it was initially tested but every other HLA-DR allele MARIA is able to predict (selection of alleles in **Fig. 1f**, full data in **Supp. Table S2**).

Mutations within linker peptides that don’t directly contribute to a protein’s activity are *a priori* less likely to reduce that activity than mutations within functional domains. To demonstrate our deimmunization workflow in the more stringent case of a direct, rather than linker-mediated, fusion of domains we applied it to the human-derived RNA-binding domain PUF-9R.^52^ PUF-9R was developed by transposing the last three repeated RNA-binding domains from the human PUM1 protein onto itself, replacing the fourth and fifth of such repeats, to create a domain that binds to a different, longer RNA sequence (**Fig. 2a**). This transposition creates two junctions not found in the original protein, between repeats 3 and 6 (the third and fourth repeats in the new protein) and repeats 8 and 6 (the sixth and seventh repeats in the new protein). The latter junction contains 15mer peptides exceeding the MARIA false omission rate threshold, which we eliminated using two mutations, V241D and H243M, at the C-terminus of repeat 8 (**Fig. 2b**). The other junction, between the native repeats 3 and 6, had no peptides above the MARA threshold so no mutations were required. We then assayed both the original and deimmunized PUF-9R domains by fusing them to the human ADAR2 deaminase domain (ADAR2dd) and co-transfecting them with a plasmid encoding a reporter transcript. The GFP output of this reporter is dependent on the deamination of a stop codon (UAG) to a non-stop codon (UIG), which will more readily occur when the ADAR2dd is colocalized to the stop codon by PUF-9R’s binding to a nearby site in the reporter transcript (**Fig. 2c**).^53^ We found that introducing the deimmunizing mutations reduced but did not eliminate the fluorescent output (**Fig. 2f**, “V241D, H243M”), likely because these mutations occur in a structured portion of the PUF repeat and disrupt its stability or target binding.

**Fig. 2.**
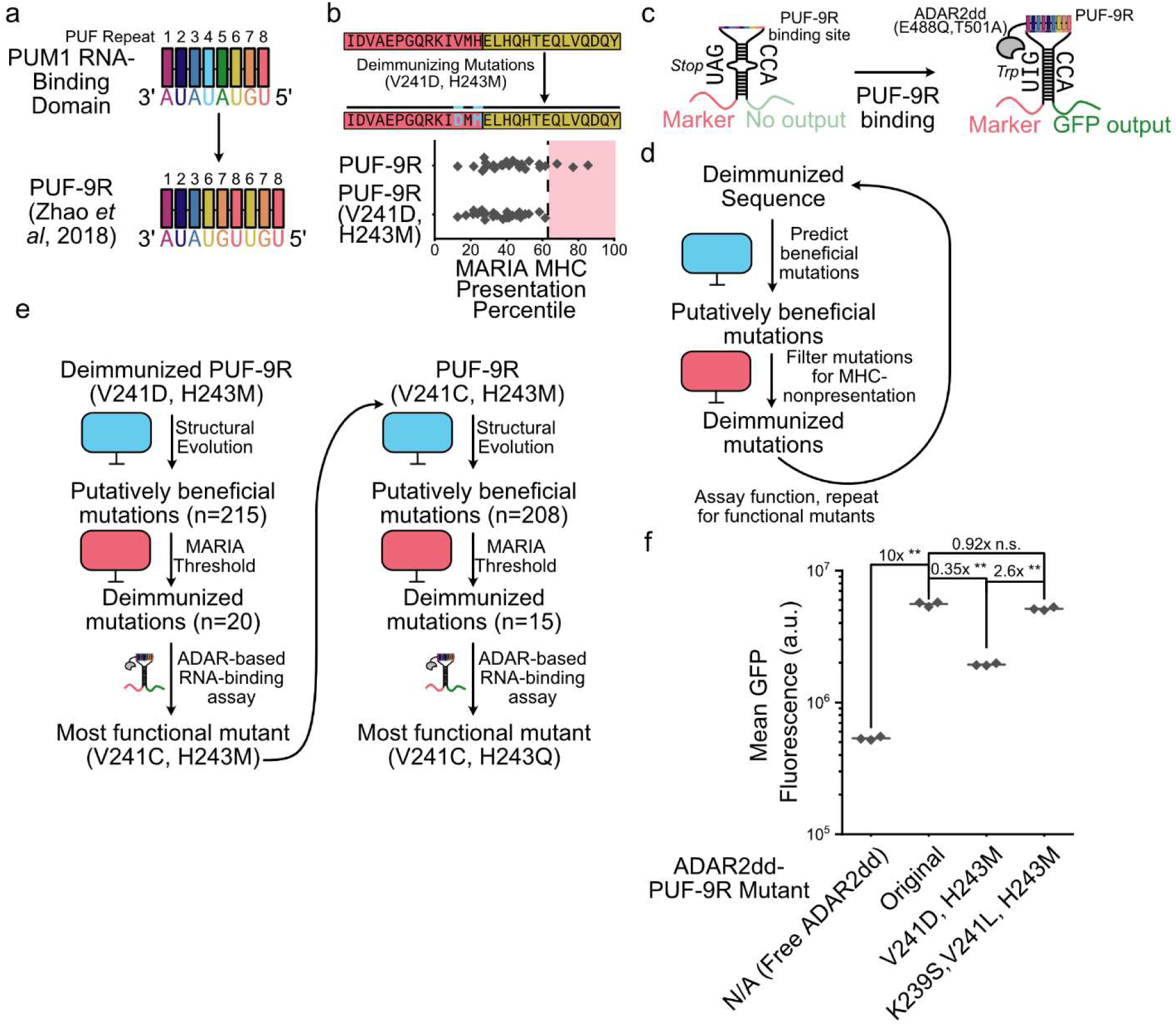
Deimmunization and optimization of PUF-9R RNA-binding domain. (**a**) Schematic of PUF-9R domain, showing its derivation from the human PUM1 RNA-binding domain. (**b**) Deimmunization of the PUF repeat 3-PUF repeat 1 junction in PUF-9R. 15mer peptide identities and MARIA percentile scores are in **Supp. Table S5**. (**c**) Schematic of ADAR-based RNA-binding assay. (**d)** Schematic of immunogenicity-sensitive mutational optimization workflow. (**e**) Mutational optimization applied to PUF-9R (V241D, H243M). (**f**) Flow cytometric ADAR-based RNA binding assay for PUF-9R variants transfected along with reporter plasmid. ** denotes p < 0.005 and n.s. denotes p > 0.05.

This test case offers an opportunity for us to introduce an additional algorithm to address a common challenge that we anticipate: the need to improve the function of engineered proteins. Due to the deimmunization constraint, conventional directed evolution, which generates large numbers of mutated sequences that cannot easily be limited to ones which are not immunogenic, isn’t suitable. Instead, we employed a secondary workflow of immunogenicity-sensitive mutational optimization (**Fig. 2d**). Using a structure-informed protein evolution method developed by Shanker *et al*,^54^ we identified mutations in the deimmunized protein sequence which would increase the sequence’s likelihood of folding into its own structure according to the protein language model ESM-1F. The structure of the deimmunized protein was predicted using the AlphaFold 3 server.^55^ Like other algorithmic choices in this workflow, ESM could be replaced by other protein language models^56–64^ for determining beneficial mutations. We then filtered these mutations based on the MARIA score of the newly generated nonhuman 15mers, created plasmid constructs with the deimmunized mutations, and assayed them to determine which mutations increased the function of the RNA-binding protein. We then repeated the workflow on the more-functional mutant sequences to find an optimal sequence subject to the deimmunization constraint. For the deimmunized PUF-9R domain, two rounds of this process were performed before reaching highly diminishing returns from additional mutations (**Fig. 2e**). Our final construct, with the mutations V241C, and H243Q, restored the function of PUF-9R to a level comparable to the original domain while deimmunizing all of its nonhuman peptides (**Fig. 2f**).

As a more complex subject of deimmunization, we turned to DNA-binding domains, which as described above generally require novel, nonhuman protein sequences to bind to novel DNA sites. The clearest way to circumvent this issue is to combine portions of human DNA-binding domains such that they bind a new motif that is a concatenation of the sub-motifs bound by the source domains. This strategy has proven effective with arrays of human zinc fingers^65^ or other DNA-binding domains such as homeoboxes.^66^ We seek to expand on this method, however, by using our workflow to ensure that the junctions between the component domains, the only nonhuman parts within them, are deimmunized. Mutational deimmunization as performed with NZF and PUF-9R may pose problems with ZFs, however, as the linkers between ZFs are strongly conserved, with mutations in them weakening interactions between the ZF array and the DNA backbone.^67^ The small size of zinc finger domains also means that deimmunizing mutations could risk disrupting either the DNA-binding residues or the alpha helix that contains them. Because of this, we sought non-natural junctions between human ZFs which would already, without introducing mutations, have no 15mer peptides above the MARIA threshold.

We started by computationally screening for ZF fusions which bound non-genomic sites, echoing the function of DNA-binding domains like rtTA. We identified a zinc finger array that fuses a four-finger block from ZNF35 to one of the same size from ZNF250 (**Fig. 3a**) wherein the junction between these two blocks would produce no nonhuman peptides above the MARIA threshold (**Fig. 3d**, “ZNF35/ZNF250”). We predicted this composite domain would selectively bind to a composite motif made by combining the two blocks’ respective target motifs (**Supp. Fig. S1**). The full “ZNF35/ZNF250” binding domain induces transcription of a co-transfected GFP reporter when the binding domain is fused to the previously described NZF transcriptional activator and the GFP is downstream of the predicted binding site (**Fig. 3b**, replicate data in **Supp. Fig. S1**). Moreover, whenever the ZNF35/ZNF250 binding domain was replaced with only the portion derived from one of its source proteins, or whenever the corresponding binding site was replaced with that predicted to bind to only one of the source proteins, the GFP production was eliminated. This indicates that the engagement of both sub-domains is necessary for transcriptional activation by this synthetic TF.

**Fig. 3.**
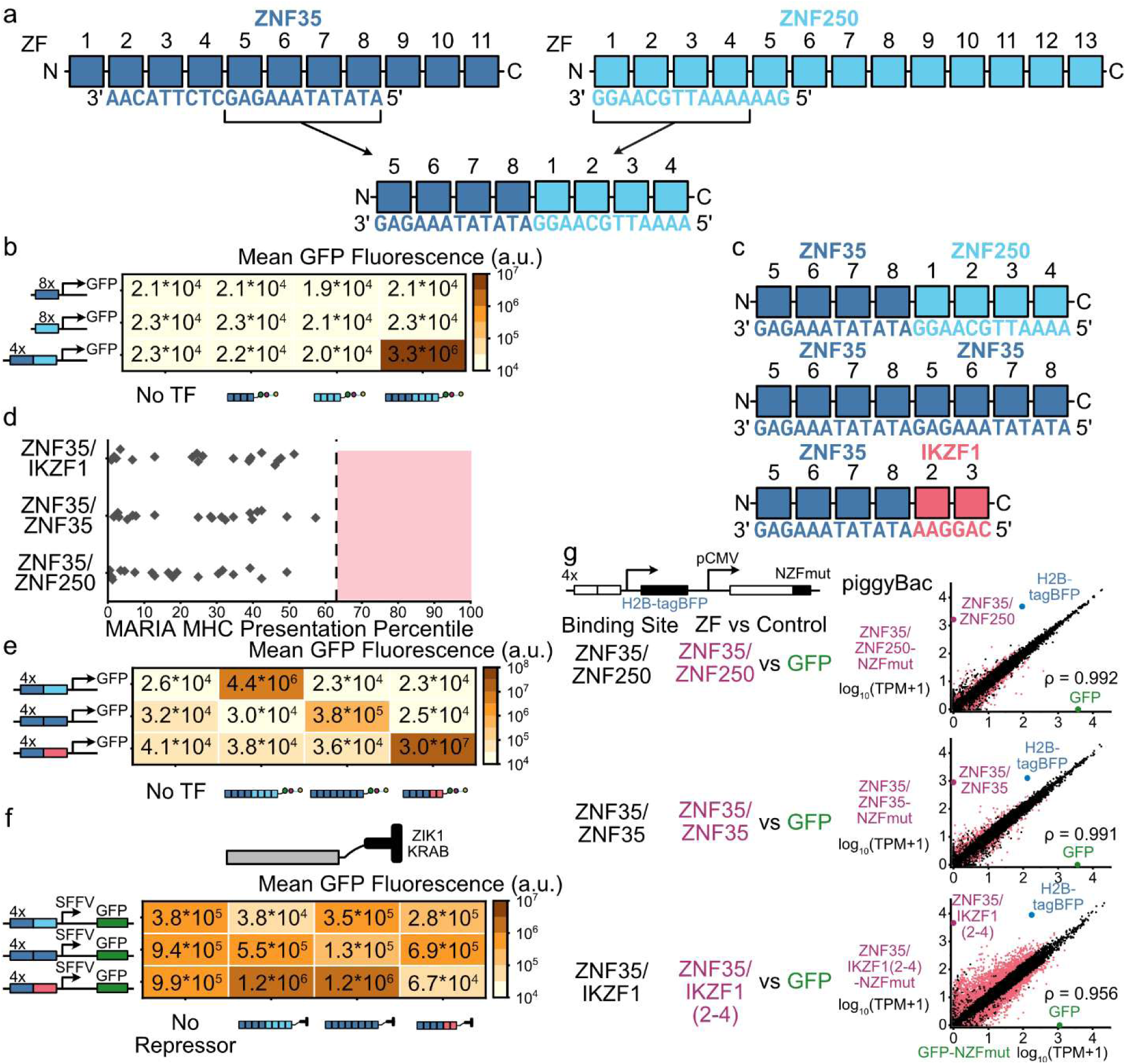
Deimmunized block fusion zinc finger arrays for genome-orthogonal DNA binding. (**a**) Construction of ZNF35/ZNF250 DNA-binding domain. (**b**) Flow cytometry results from transfection of plasmids encoding ZNF35/ZNF250-NZF transcription factor and a reporter containing four copies of its binding site upstream of a GFP reporter, or combinations of the two containing only the portions derived from either ZNF35 or ZNF250. In all heatmaps unless otherwise noted, values represent the mean of three biological replicates. Replicate data are in **Supp. Fig. S1**. (**c**) Schematic of three designed block fusion DNA-binding domains, ZNF35/ZNF250, ZNF35/ZNF35, and ZNF35/IKZF1, with the DNA sequences to which they are predicted to bind. Amino acid sequences for zinc finger arrays are in **Supp. Table S1.** (**d**) MARIA scores of junction peptides in block fusion zinc finger arrays. Points represent the MARIA percentiles for all 15mer peptides containing amino acids from both human proteins for the HLA-DRB1*01:01 allele. 15mer peptide identities and MARIA percentile scores are in **Supp. Table S5**. (**e**) Flow cytometry results from transfection of HEK293 cells with plasmids encoding transcription factors containing each block fusion zinc finger array joined to the NZF transcriptional activation domain along with reporters containing four copies of their corresponding binding sites upstream of a GFP reporter. Replicate data are in **Supp. Fig. S2a**. (**f**) Flow cytometry results from transfection of HEK293 cells with plasmids encoding transcriptional repressors containing each block fusion zinc finger array joined to the ZIK1 KRAB domains domain along with reporters containing four copies of their corresponding binding sites upstream of an SFFV promoter-driven GFP reporter. Replicate data are in **Supp. Fig. S2b**. (**g**) Transcriptomic analysis of FLP-IN T-REx-293 cell lines with piggy-Bac-integrated plasmids encoding block fusion zinc finger transcription factors or GFP-NZF controls and corresponding H2B-tagBFP reporters. ρ denotes the Pearson correlation coefficient between the transcript per million (TPM) values of the two conditions. Red points on scatter plots denote transcripts for which the fold change between the two conditions is greater than 2 and the adjusted p value calculated by DESeq2 (see Methods) is less than 0.05.

To expand on this single deimmunized, putatively genome-orthogonal DNA-binding domain, we generated two additional ones via the same procedure, each of which contained the four zinc fingers from ZNF35 fused to either itself (“ZNF35/ZNF35”), or two zinc fingers from IKZF1 (“ZNF35/IKZF1”) (**Fig. 3c**). All the nonhuman junction peptides in these constructs fall below the MARIA false omission rate threshold (**Fig. 3d**). When co-transfected with GFP reporter plasmids containing four copies of their binding sites, all three of the fusion zinc fingers were able to increase GFP expression, while exhibiting no off-target activation between different ZF-binding site pairs (**Fig. 3e**, replicate data in **Supp. Fig. S2a**). Additionally, each of these transcription factors requires the binding of both subdomains derived from each of the proteins to function (**Supp. Fig. S3**). Replacing the NZF activation domain with a human transcriptional repressor, the KRAB domain from ZIK1, shows that these fusion ZFs can repress a constitutively active SFFV promoter with adjacent binding sites (**Fig. 3f**; replicate data in **Supp. Fig S2b**). The off-target effects for transcriptional repression were more significant than for activation, possibly because more transient interactions between the common ZNF35 binding subdomain and site were able to induce DNA methylation and promoter repression by the KRAB while more sustained binding is necessary to maintain transcription.

To test to what extent our fusion ZFs had off-target genomic effects we used the piggyBac transposon to create stable cell lines that expressed them and performed RNA-seq to find transcripts that were differentially expressed between those lines and ones with GFP integrated as a control (**Fig. 3g**; full analysis in **Supp. Fig. S4**). The results are variable, with the ZNF35/IKZF1 integrant in particular (which, it is worth noting, is an earlier version of this construct containing IKZF1 ZFs 2 to 4 rather than 2 and 3, thus having seven rather than six ZFs total) having widespread off-target effects, in contrast to the modest off-target effects for the other two. This could be an artifact of the potentially unequal number of plasmids integrated by piggyBac or different expression of the ZFs in different cell lines, as the number of ZNF35/IKZF1 transcripts per million was greater than that of the other integrated ZFs. Conversely, it could indicate actual increased off-target binding and activation by the ZNF35/IKZF1 TF than the other two.

A key benefit of synthetic transcription factors is that they can act as “tuning knobs” activated or deactivated by small molecule drugs. Therefore, incorporating that ability is desirable, although adding one or more new domains for external control will introduce new junctions that must be deimmunized. One way to get around this is to incorporate control directly into the DNA-binding domain. ZF 2 of IKZF1, which is included in our ZNF35/IKZF1 binding domain, is targeted for degradation by immunomodulatory drugs (IMiDs), such as pomalidomide (Pom).^68,69^ This zinc finger has been used by itself as a degron,^70–72^ but since it is already present as part of the DNA-binding domain, pomalidomide should be able to induce the degradation of ZNF35/IKZF1 (**Fig. 4a**). Indeed, adding pomalidomide to HEK293 cells co-transfected with a ZNF35/IKZF1 transcriptional activator and the corresponding GFP reporter reduced GFP fluorescence, while it had no effect on ZNF35/ZNF250-induced GFP (**Fig. 4b**). A more conventional control method would be the rapalog (Rap)-induced association of the DNA-binding and transcriptional activation domains via their fusion to the mTOR FKBP-rapamycin binding (FRB) domain and FKBP itself.^73^ This is enabled in our workflow by deimmunizing the linkers between the functional and rapalog-binding domains (**Fig. 4c**, deimmunization details in **Supp. Fig. S5**). However, the T2098L mutation in FRB, which causes it to bind to the biorthogonal rapamycin analogue AP21967 as well as rapamycin itself, introduces nonhuman peptides above the MARIA false omission rate threshold. Introducing a second mutation, N2093W, resolves this issue (**Fig. 4d**) without substantially impacting rapalog-dependent transcriptional activation (**Supp. Fig. S5)**. Co-transfecting HEK293 cells with NZF attached to the double mutant FRB, ZNF35/ZNF250 attached to FKBP, and the ZNF35/ZNF250-sensitive GFP reporter plasmid shows GFP activation only when rapalog is added, indicating that the deimmunization of additional linkers and the FRB domain is compatible with rapalog-dependent gene activation (**Fig. 4e**).

**Fig. 4.**
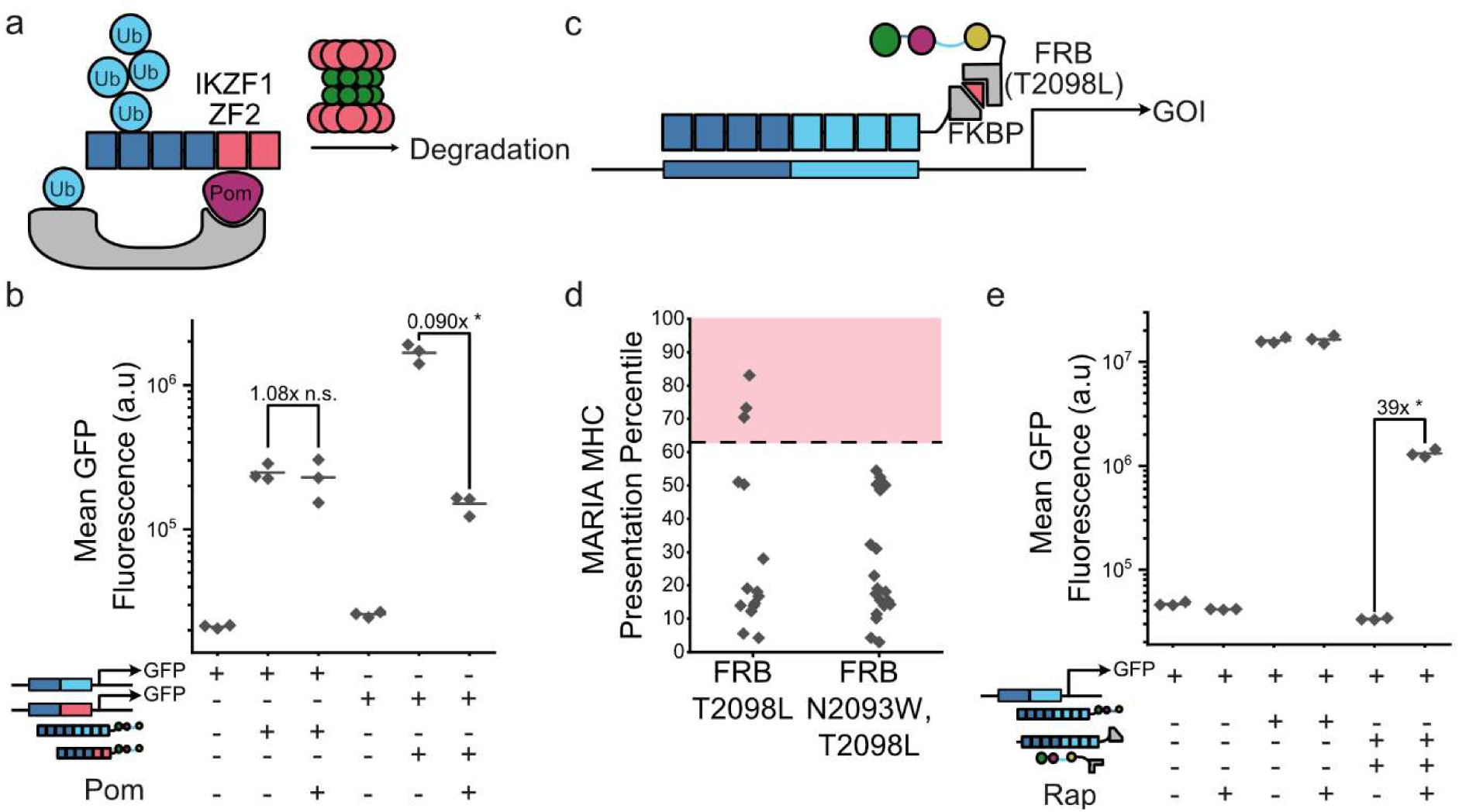
Small molecule-based control of deimmunized block fusion zinc finger transcription factors. (**a**) Schematic of pomalidomide-based degradation of the ZNF35/IKZF1 DNA-binding domain. (**b**) Flow cytometric measurement of selective control of ZNF35/IKZF1-containing transcription factors using pomalidomide. * denotes p < 0.05 and n.s. denotes p > 0.05. “+ Pom” conditions received 1 µM Pom; “-Pom” conditions received 1/1000 DMSO vehicle. (**c**) Schematic of rapalog-induced activation of a block fusion zinc finger transcription factor. (**d**) Deimmunization of rapalog-specific FRB domain. 15mer peptide identities and MARIA percentile scores are in **Supp. Table S5**. (**e**) Flow cytometric measurement of the control of split ZNF35/ZNF250 transcription factor using rapalog-induced dimerization. * denotes p < 0.05. “+ Rap” conditions received 500 nM rapalog; “-Rap” conditions received 1/1000 ethanol vehicle.

The “block fusion” zinc finger arrays described above have value in controlling artificially introduced transgenes. The complement of this strategy is to generate DNA-binding domains that can target specific genomic sites like Cas proteins can. Building on previous examples,^65^ we aim to ensure the deimmunization of nonhuman peptides at the ZF-ZF junctions. We also seek to leverage advances in the prediction of ZF binding specificities^74–78^ to increase the number of distinct ZFs that can be used in these domains without needing to validate their binding specificities individually. The workflow we have established is shown in **Fig. 5a**. First, the entire set of human zinc fingers is filtered by the requirement that specific linker sequences must be used to join ZFs together, and that the addition of these sequences must not introduce potentially immunogenic peptides. Therefore, we appended a linker peptide with sequence TGERP to the N and C termini of every human ZF and accepted only those ZFs where this introduced no 15mer peptides above the MARIA false omission rate threshold of about the 63^rd^ percentile. Of the 6892 distinct ZFs identified in the human genome, 947 (13.7%) are below this MARIA threshold, while when the same percentile threshold is applied with the NetMHCIIpan algorithm,^34^ 1133 (16.4%) pass it (**Fig. 5b**). While these values are comparable, only about one fifth of the ZFs accepted by each algorithm are accepted by the other, highlighting that the choice of algorithm has a significant role. Future, more accurate tools for predicting MHC presentation will enable our modular workflow to more consistently produce bona-fide non-immunogenic constructs.

**Fig. 5.**
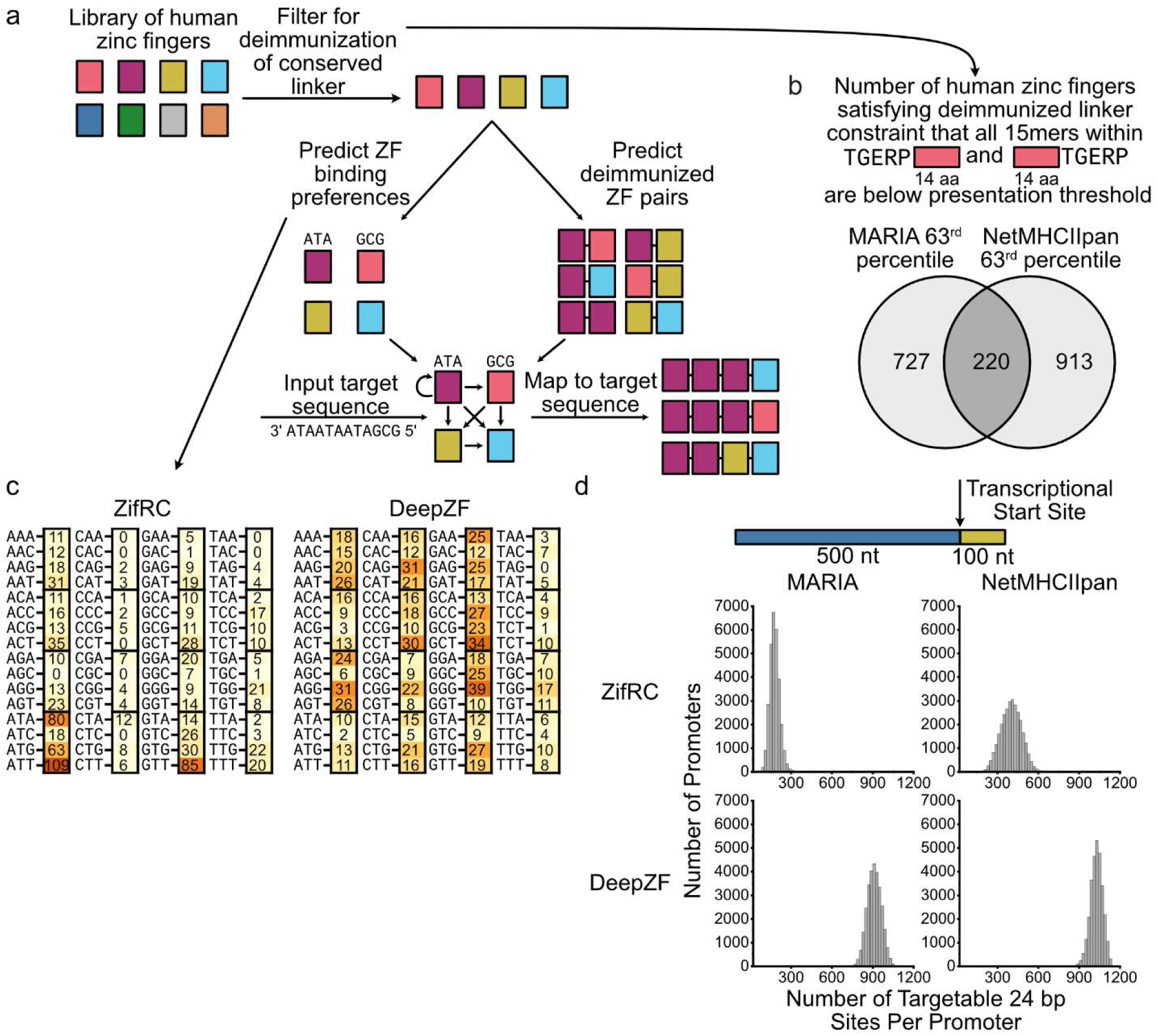
Genome-targeting deimmunized zinc finger construction workflow and comparison of algorithms for use in that workflow. (**a**) Schematic of workflow for constructing deimmunized genome-targeting zinc finger arrays. (**b**) Number of ZFs with satisfying deimmunized linker constraint for MARIA and NetMHCIIpan MHC-presentation prediction algorithms. (**c**) Number of ZFs that satisfy deimmunized linker constraint for MARIA which bind to each trinucleotide according to ZifRC and DeepZF zinc finger affinity-prediction algorithms. (**d**) Number of targetable 24 bp sites for every human promoter according to each combination of MHC prediction and ZF affinity algorithms.

We then computationally predicted the trinucleotide binding specificity for each ZF in the acceptable set. We determined the position weight matrix for each ZF and took the nucleotide for each position with the maximum weight. Again, the choice of algorithm for predicting MHC binding is significant. For example, ZifRC^74^ predicts that the ZFs accepted by MARIA are biased towards a handful of trinucleotides, while DeepZF^77^ predicts a more even distribution (**Fig. 5c**). In parallel, the junction sequences from every pair of ZFs in the acceptable set were analyzed with the MHC-presentation predictor to find every ZF-ZF junction that is not predicted to be presented by the MHC-II allele in question. The accepted ZF pairs define those that can be used in succession in the constructs we design. This means that, to bind a DNA motif, we first take the trinucleotide at the 3’ end of this sequence (where the most N-terminal ZF will bind) and find all the accepted ZFs which bind to it. For each of those ZFs, we then determine which ZFs both bind to the next trinucleotide and can be placed after the first ZF in an accepted pair. We can repeat this process to get all the possible arrays of ZFs for a target sequence.

We determined the number of 24 bp sites within every human promoter in EPDnew^79^ (defined as the region from 500 bp upstream of each transcriptional start site to 100 bp downstream of it) which could be bound by at least one ZF array created using our method. We did this with each combination of the two previously mentioned algorithms for predicting MHC presentation (MARIA and NetMHCIIpan) and ZF affinity (ZifRC and DeepZF) to see how algorithmic choice affected the restrictions on ZF targeting (**Fig. 5d**). The binding site size of 24 bp was chosen by analogy with the previously described block ZF fusion domains, and to ensure that every site chosen is likely to be unique within the human genome. We also expected that a larger DNA-binding domain would be required compared to those generated by other methods^65,80^ because the human ZFs in our constructs are chosen based on their non-immunogenicity and sequence specificity, rather than individual binding strength. Every promoter has at least some accessible sites with every pair of algorithms (the global minimum being 55) but generally using NetMHCIIpan resulted in a greater number of predicted accessible sites than MARIA, likely due to its greater number of accepted ZFs. Using DeepZF resulted in a greater number of predicted accessible sites than ZifRC, likely because it predicts fewer trinucleotides that no ZFs can bind to and a generally more even distribution of predicted ZF affinities. These different distributions should not be interpreted as implying that one algorithm is “better” than another: the important factor is how good each algorithm is at its own function. The ground truth of truly non-presented junction peptides and ZF affinities may be relatively restrictive or relatively loose and can only be determined or approximated as each type of algorithm becomes more accurate. To show that even the more restrictive choices can still permit the design of functional ZF arrays, we opted to use MARIA and ZifRC, the most restrictive pair of algorithms, to create genome-targeting transcription factors.

To demonstrate our process for generating deimmunized genome-targeting ZF arrays we targeted the genes utrophin (UTRN) and sodium voltage-gated channel alpha subunit 1 (SCN1A). Overexpression of each of these genes shows potential therapeutic promise for particular diseases. UTRN is a homolog of dystrophin, meaning that expression of UTRN in skeletal muscle could relieve Duchenne muscular dystrophy.^81,82^ Haploinsufficiency of SCN1A, meanwhile, causes Dravet syndrome, an epileptic encephalopathy, and increasing the production of the functional SCN1A copy may be able to relieve it.^83,84^ However, the coding sequences of both UTRN and SCN1A exceed the roughly 4 to 5 kb delivery limit of adeno-associated viruses,^85^ making direct delivery of those genes more impractical than the introduction of transcription factors that increase expression of their genomic copies. Using the algorithm described above (**Fig. 5**) with MARIA as the MHC-presentation predictor and ZifRC as the ZF affinity predictor, we found that there were 123 and 130 targetable 24 bp sites in the promoters of UTRN and SCN1A, respectively (**Fig. 6a**). For each of these sites we found every possible human-derived ZF array with deimmunized ZF-ZF junctions that could bind to them. We determined the binding score—defined as the sum of the position weight matrix values for each nucleotide in the predicted binding motif—of every such array, allowing us to find the highest-scoring ZF array for each site (**Fig. 6b**).

**Fig. 6.**
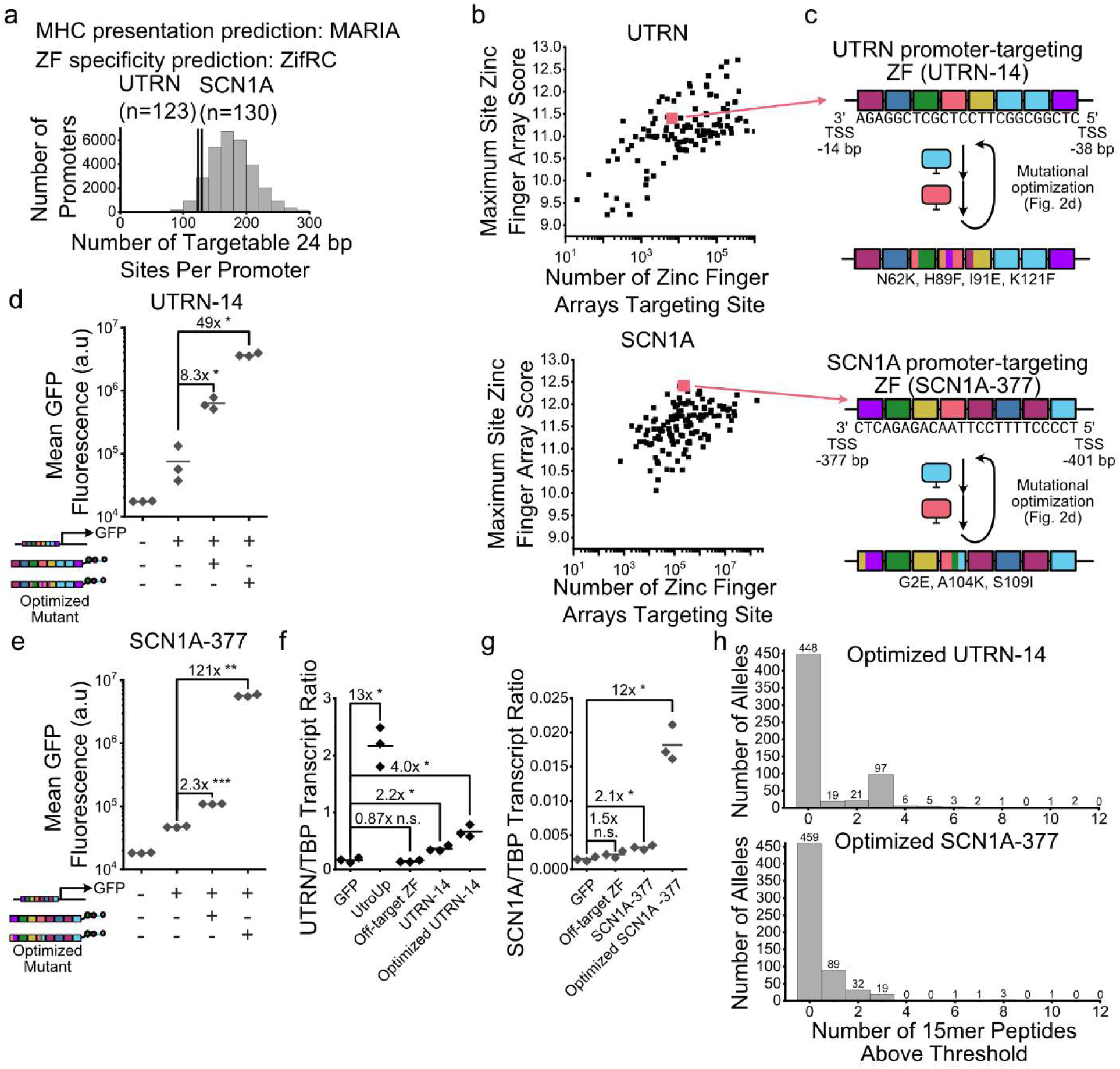
Generation of deimmunized zinc finger arrays targeting UTRN and SCN1A. (**a**) Number of targetable sites in UTRN and SCN1A promoters. (**b**) Number of sites and maximum array score for all targetable sites in UTRN and SCN1A promoters. (**c**) Selected zinc finger arrays targeting UTRN and SCN1A, their corresponding binding sites, and mutations present in their optimized sequences. Amino acid sequences for zinc finger arrays are in **Supp. Table S1.** (**d**) Flow cytometric measurement of transcriptional activation of a co-transfected GFP reporter from UTRN-14 transcription factor. * denotes p < 0.05. (**e**) Flow cytometric measurement of transcriptional activation of a co-transfected GFP reporter from SCN1A-377 transcription factor. ** denotes p < 0.005 and *** denotes p < 0.0005. (**f**) Reverse-transcription quantitative PCR measurement of UTRN transcript upregulation due to UTRN-14 and other transcription factors. * denotes p < 0.05 and n.s. denotes p > 0.05. (**g**) Reverse-transcription quantitative PCR measurement of SCN1A transcript upregulation due to SCN1A-377 and other transcription factors. * denotes p < 0.05 and n.s. denotes p > 0.05. (**h**) Number of 15mer peptides from the optimized UTRN-14 and SCN1A-377 transcription factors above MARIA threshold for all HLA-DR alleles that can be predicted by MARIA.

We selected one ZF array and corresponding site as a final candidate for each of UTRN and SCN1A (**Fig. 6b**). The UTRN array binds a site 14 bp upstream of its transcriptional start site and was selected for being located adjacent to another targetable site, potentially allowing cooperative binding^24^, though the ZF ostensibly targeting the adjacent site was not able to activate transcription (**Supp. Fig. S6)**. The SCN1A array binds a site 377 bp upstream of its transcriptional start site and was selected for having the highest binding score across all sites and having the greatest activation among the top 10 scoring non-redundant SCN1A-targeting arrays (details regarding non-selected ZF arrays are in **Supp. Fig. S6**; ZF array sequences are in **Supp. Table S1**). Owing to the location of their binding sites relative to the respective transcription start sites the ZFs were termed UTRN-14 and SCN1A-377 respectively. UTRN-14 was subjected to four rounds of immunogenicity-filtered mutational optimization as described above (**Fig. 2d, e**) while SCN1A-377 was subjected to three. The identification of probability-increasing mutations in the first three rounds of evolution for UTRN-14 was done with an earlier structure-insensitive evolution algorithm,^86^ while for the last we used the structure-sensitive algorithm as in **Fig. 2**.^54^ The first round of SCN1A-377 mutation was done with the structure-insensitive algorithm while the latter two used the structure-sensitive algorithm.

Both UTRN-14 and SCN1A-377, when fused to NZF, induced transcription of a fluorescent reporter when co-transfected with reporter plasmids containing a single copy of their predicted binding sites directly upstream of GFP (**Fig. 6d, e**), with the evolved mutants having a greater ability to induce transcription in both cases. Similarly, both sets of ZF arrays induced a significant increase in the transcript levels of their targeted genes—UTRN or SCN1A—when transfected into HEK293 cells, indicating that they were able to bind to the genomic locus of those genes and induce their transcription (**Fig 6f, g**). Significantly, though, transfection of UTRN-14 resulted in a lower level of UTRN production than UtroUp, a previously published UTRN-targeting ZF array made from nonhuman ZFs^87^. UTRN-14 was optimized using the less-sophisticated structure-insensitive algorithm, however, while SCN1A-377, which was optimized partially with the structure-sensitive algorithm, had a substantially greater fold change. This indicates that better methods of protein optimization will likely allow human-derived ZF arrays to meet or exceed the function of ones derived from directed evolution. UTRN-14 was also not the highest-scoring ZF across all UTRN promoter sites, while SCN1A-377 was (**Fig. 6b**). To see how portable our ZF arrays were to other MHC-II alleles besides the one for which they were designed we scored all the nonhuman peptides in those arrays using every HLA-DR allele MARIA can predict. We found that for both UTRN-14 and SCN1A-377 no 15mer peptides were above the MARIA false omission rate threshold for most alleles, with only two alleles predicting as many as 11 potentially immunogenic 15mers for UTRN-14 or one predicting 10 for SCN1A-377 (**Fig. 6h**). Further engineering would be needed to eliminate those immunogenic peptides to create versions of these transcription factors suitable for all patients. However, the concentration of alleles that are predicted to have no presented peptides suggests that allelic variation, at least among HLA-DR alleles, would not require patient-specific engineering of all proteins made with our workflow.

## Discussion

In this study, we have established a workflow for combining algorithms to simultaneously engineer clinically relevant proteins and maintain low immunogenicity. We identified potentially immunogenic peptides based on their predicted presentation by the MHC and deimmunized them by introducing mutations or judiciously selecting fusion domains. This approach combines two pre-existing trends in the addressing of immunogenicity, namely the prediction and deletion of T-cell epitopes within non-self proteins^88–94^ and the use of human or “human-like” domains in place of those sourced from other species, such as zinc fingers rather than TALEs or Cas proteins for DNA binding.^14^ We expect a workflow that combines these practices will create proteins that are more strictly deimmunized than either of them in isolation. The use of human-derived domains limits nonhuman peptides to small sequences around mutations and interdomain junctions, drastically reducing the number of peptides potentially requiring deimmunization compared to nonhuman proteins. This decreases the risk of false negatives in epitope prediction, which is intrinsic to imperfect prediction algorithms but can be reduced by decreasing the number of peptides with the potential for immunogenicity. Removing peptides which are presented by MHC molecules is also a stricter criterion than removing T cell epitopes—peptides which can be bound by some TCR—because MHC presentation is a precondition for TCR binding. The relative strictness of the non-immunogenicity requirements raises the question of whether functional proteins can be constructed while respecting them. We have shown, however, that mutations sufficient to deimmunize chimeric human proteins may either have no measurable effect on protein function, as with NZF (**Fig. 1**), or may be rescuable through immunogenicity-filtered optimization, as with PUF-9R (**Fig. 2**).

The first set of deimmunized zinc finger transcription factors presented in this paper are “block fusions,” fusions of continuous blocks of zinc fingers derived from human proteins, designed to be genome-orthogonal and used for controlling the transcription of transgenes. The three such block fusion ZF arrays presented above are largely orthogonal to one another, allowing their simultaneous use for independent control of different genes. There are substantial differences between their ability to induce transcription, however, with the difference between the fold activations of ZNF35/IKZF1 and ZNF35/ZNF35 being a factor of about 100. This indicates that creating more such block fusions would be desirable, so that multiple different ZF arrays with comparable strength can be used at once. It would also likely be prudent to avoid using the N-terminal ZNF35 ZF 5-8 sub-array common to all the existing block fusions, as this may reduce the residual off-target effects seen when repressing transgenes. The extent to which integration of the synthetic transcription factors influences the levels of genomic transcripts is also of concern. Part of the reason for the greater number of genomic off-targets for ZNF35/IKZF1 may be its stronger DNA binding than the other ZFs arrays we created. This is because if the binding affinity of a small portion of a ZF array, which binds to a small DNA sequence that is likely widely repeated in the genome, is strong enough, then the ZF array may bind to those sequences even if the rest of the array doesn’t have significant affinity for the surrounding DNA residues. Equalizing the binding strength of the individual ZFs may lessen that effect, since every individual ZF’s contribution will be relatively small and more of them would have to be engaged for the protein to bind substantially. This may be challenging, however, due to the diversity of ZF-DNA binding strengths^95^ and the fact that most ZF affinity predictors predict sequence specificity in the form of PWMs rather than binding strength.

Most notably, we have demonstrated the generation of deimmunized ZF arrays targeting genomic DNA sequences for use in regulating the transcription of endogenous genes. This method has a particular reliance on the accuracy of external models since as presented it relies on the prediction of both MHC-presented peptides and the DNA sequence specificity of zinc fingers. Another key challenge lies in the number of potential arrays which can target a given sequence. While some sites will have no possible deimmunized ZFs targeting them, those which do may have up to hundreds of millions, requiring experimental library validation in a library format or even beyond. Conversely, we may be able to generate more diverse or functional ZF arrays by reconsidering their structure. For example, our current method assumes that the conserved linker (TGERP) is not naturally adjacent to any ZF, and that therefore any peptide that contains any residue from that linker is nonhuman and should be analyzed for immunogenicity. This is a stricter criterion than is necessary because some ZFs will naturally have this linker at their N or C terminus, meaning that even if peptides containing only amino acids from that ZF and the linker are presented by MHC they should still be recognized as self. Recognizing this and using multiple different ZF linkers which naturally surround the ZFs used would increase the number of possible ZF arrays we can construct. This would increase the likelihood of some being more functional, at the cost of increasing the technical and computational complexity of the algorithm for generating zinc finger arrays. Nevertheless, the fact that we have generated functional activators of endogenous genes with modest effort suggests the effectiveness of our workflow.

We have presented in this paper an application of our deimmunization workflow for diverse protein types, namely transcriptional activation domains, RNA-binding domains and DNA-binding domains. In principle, however, it is applicable to any protein primarily sourced from human proteins with non-natural fusions or mutations, whether it is directly infused for therapeutic value or expressed within an endogenous or exogenous cell. An obvious application lies in synthetic receptors which combine otherwise-unrelated ligand binding and signal transduction domains, particularly those which don’t depend on the expression of nonhuman intracellular proteases such as fully human CAR-Ts, SynNotch/SNIPRs (particularly when using human rather than mouse Notch),^15^ and various LIDAR modalities.^53^ Some enzyme-based therapies may benefit as well, particularly those which engineer human proteins to either modify their substrate specificity^96,97^ or increase their catalytic efficiency.^98,99^ As more diverse and more strongly engineered protein, gene, and cell therapies are developed, adoption of this workflow to ensure robust deimmunization offers to decrease the risk of adverse anti-therapeutic immunity and increase the safety and efficacy of such therapies.

## Methods

### Plasmid cloning

Standard molecular biology techniques were used for production of transfected plasmids, including InFusion of PCR fragments with restriction-digested plasmid backbones (Takara Bio catalogue number 638948) and ligation of backbones with digested fragments from other plasmids or annealed phosphorylated oligonucleotides. A complete list of plasmids used and maps thereof is presented in **Supp. Table S3**. Plasmids are deposited in Addgene and are available upon request from the corresponding author.

### Tissue culture

FLP-IN T-Rex HEK293 cells (Thermo Scientific catalogue number R78007) were used for generation of stably integrated cell lines (Figs 3g, S5) and standard HEK293 cells (ATCC catalogue number CRL-1573) were used for all other *in cellulo* experiments. All cells were cultured in a humidity-controlled incubator at 37 °C with 5% CO_2_. Culture media for all cells consisted of high-glucose, high-glutamine Dulbecco’s Modified Eagle’s Medium (Fisher Scientific catalogue number 501015428) supplemented with 10% fetal bovine serum (Fisher Scientific catalogue number FB12999102), 1 mM sodium pyruvate (EMD Millipore catalogue number TMS-005-C), 1× penicillin– streptomycin (Genesee catalogue number 25-512), and 1× MEM non-essential amino acids (Genesee catalogue number 25-536). Cells were tested for mycoplasma contamination and found negative.

### Transient transfection

HEK293 cells were plated in tissue culture-treated 24 well plates and cultured under standard conditions to 70-90% confluency prior to transfection. Transient plasmid transfection was performed using the jetOPTIMUS DNA transfection reagent (Polyplus catalogue number 117-15) according to the manufacturer’s recommendations, such that 0.375 µL of reagent was added to 500 ng of plasmid in 50 µL of jetOPTIMUS buffer to transfect each well of a 24 well plate. Details of plasmids used in all transfections are present in **Supp. Table S4**. In cases where additional chemicals were added to the media (Figs. 4b, 4e, S6) this was done when the media was replaced immediately before transfection. For all experiments the transfected cells were incubated for about 48 hours before further manipulation (either flow cytometry or RNA extraction).

### Flow cytometry and data analysish

Cells were trypsinized and resuspended in HBSS with 2.5 mg/mL bovine serum albumin, strained with a 40 µm filter, and analyzed by flow cytometry using a BioRad ZE5 Cell Analyzer. Flow cytometry data was analyzed using the Cytoflow Python package. Events were gated along forward and side scattering to include cells and exclude other debris and, except for Figure S5e, cells were gated for high mCherry co-transfection marker fluorescence by determining the 99.5^th^ to 99.9^th^ percentile mCherry fluorescence levels for the least transfected sample within each display item and applying that as a gate for all other samples within that display item. The geometric mean of the fluorescent output (either tagBFP or GFP depending on the experiment) was then determined for the cells within the gated population. All experiments included three biological replicates for each condition. Differences between conditions were assessed for significance using a Bonferroni-corrected two-tailed Welch’s *t-*test. Statistical analyses and graph generation were done in OriginPro.

### Cell line generation

Polyclonal cell lines were generated by piggyBac integration into FLP-IN T-Rex HEK293 cells. Cells in a 24 well plate were transiently transfected with 450 ng of plasmids containing piggyBac repeats encoding a hygromycin resistance gene and 50 ng of plasmids encoding a CMV-driven piggyBac transposase. Full details of transfections can be found in **Supp. Table S4**. 48 hours after transfection the media was replaced with media containing 100 µg/mL hygromycin and hygromycin-resistant cells were selected for over 14 days. All cells that survived hygromycin selection were collected and pooled. The polyclonal lines were characterized by seeding them into 24 well plates, incubating them with media either containing or lacking 100 ng/mL doxycycline to induce transcription of the CMV-driven transcription factor or GFP-NZF control, and performing flow cytometry after 48 hours (Fig. S5e). For RNA-seq analysis, the polyclonal cell lines were incubated with media containing 100 ng/mL doxycycline for 48 hours before RNA extraction.

### RNA extraction

RNA for RNA-seq and reverse-transcription quantitative PCR was extracted from cells grown in 24 well plates using the RNeasy Mini QIAcube Kit (Qiagen catalogue number 74116) with cell disruption using QIAshredder columns (Qiagen catalogue number 79656) and genomic DNA digestion with the Qiagen RNase-Free DNase Set (Qiagen catalogue number 79254).

### RNA-seq and transcriptomic analysis

RNA-seq was performed on an Illumina NovaSeq X Plus by Novogene (Sacramento, CA). 100 ng total RNA was provided for two biological replicates from each cell line, from which mRNA was obtained by poly-A enrichment. At least 44 million 150 bp paired-end reads were obtained from each sample. Quality control of reads was performed using fastp^100^ with default settings for paired-end reads. Filtered reads were mapped to the hg38 reference genome using HISAT2^101^ and the mapped reads were summarized to determine the number of reads per transcript using featureCounts.^102^ Determination of differentially-expressed genes between conditions was performed using the DESeq2 R package.^103^

### Reverse-transcription quantitative PCR (RT-qPCR)

cDNA libraries were generated from extracted RNA using the iScript cDNA synthesis kit (Bio-Rad catalogue number 1708891). Quantitative PCR was performed using a QuantStudio 6 Pro (Applied Biosystems) with the SsoAdvanced Universal SYBR Green Supermix kit (Bio-Rad catalogue number 1725271). The following quantitative PCR primers were used for each targeted gene:

### TBP

> Forward: TGTATCCACAGTGAATCTTGGTTG

> Reverse: GGTTCGTGGCTCTCTTATCCTC

### UTRN

> Forward: CAAACACCCTCGACTTGGTT

> Reverse: TGGTGGAGCTGCTATCAGTG

### SCN1A

> Forward: GGACTGTATGGAGGTTGCTGGT

> Reverse: GCAAGGTTGTCTGCACTAAATGAG

### Human zinc finger identification

To find every human protein containing a C2H2 zinc finger domain, a Uniprot search was performed with the parameters “(organism_id:9606) AND (ft_zn_fing:c2h2)”. To find the locations and sequences of zinc fingers in the resultant proteins without relying on annotations which could be inconsistent in their boundaries (such as by including or excluding adjacent linkers, for example), a regular expression searching for strings of the form XXCXX[XX]CXXXXXXXXXXXXHXXX[XXXX][HC]—that is, two arbitrary residues, followed by two cysteines separated by two to four arbitrary residues, followed by exactly twelve residues, followed by either two histidines or a histidine followed by a cysteine separated by three to seven arbitrary residues—was applied to all the protein sequences and the nonoverlapping, nonredundant matches were collected as the complete set of human C2H2 zinc fingers.

### MHC presentation prediction and threshold determination

Prediction of MHC presentation with MARIA^41^ was performed using the MARIA webserver (https://maria.stanford.edu/). Unless stated otherwise, predictions were performed using the HLA-DRB1*01:01 allele. All peptide sequences analyzed were 15 amino acids in length. Determination of the false omission rate for different MARIA percentile thresholds was performed using representative receiver operating characteristic (ROC) curve data obtained via personal communication with the MARIA authors. The ROC curve points, which correspond to values of the model’s true positive rate (TPR) and false positive rate (FPR) for different percentile cutoffs were used to calculate the false omission rate (FOR) for those cutoffs with the equation

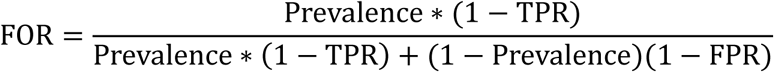

where the prevalence, the fraction of peptides actually presented by MHC, was taken as 1% (0.01). Prediction of MHC presentation with NetMHCIIpan^34^ was performed locally using NetMHCIIpan 4.2. All analyses with NetMHCIIpan used the HLA-DRB1*01:01 allele and the peptide input was the conserved ZF linker with 14 amino acids from the surrounding ZFs at either end of the linker.

### Zinc finger binding specificity prediction

Prediction of zinc finger binding specificity with both ZifRC^74^ and DeepZF^77^ were performed locally. For ZifRC the input was each individual finger, without adjacent linker sequences. For DeepZF the input was the 12 helix residues in each finger and the provided model was used without additional fine-tuning.

## Data Availability

Plasmids and plasmid maps will be deposited to Addgene prior to final publication. RNA-seq data is deposited to the Sequence Read Archive with accession number PRJNA1198109. Full flow cytometry data is available upon request from the corresponding author.

## Code Availability

Code for generating human-derived non-immunogenic zinc finger arrays is available at https://github.com/ericwolfsberg/deimmunized-human-derived-zf.

## Acknowledgements

This research was supported by NIH (R00EB027723, DP2OD034951; X.J.G), Longevity Impetus Grants (X.J.G.), Stanford ChEM-H Seed Grant (X.J.G., A.A.A.), Stanford Bio-X Interdisciplinary Initiatives Seed Grant Program (IIP) [R11-7] (X.J.G., A.A.A.)

## Authorial Contributions

E.W. and X.J.G. conceived of and directed the study. J.S.P. designed, optimized, and tested all SCN1A-targeting zinc fingers. B.C. and A.A.A. provided ROC data from and advice on the use of MARIA. J.T., L.B., and M.C.B. provided pre-publication access to the NZF transcriptional activation domain. E.W and J.S.P wrote the code for designing zinc fingers targeting genomic sites. All other experiments were developed by E.W. and X.J.G. and performed by and analyzed by E.W. E.W. and X.J.G wrote the manuscript with input and feedback from all authors.

