## Supplemental Figures S1-S6 and Supplmental Table S1 for "Machine-Guided Dual-Objective Protein Engineering for Deimmunization and Therapeutic Functions"

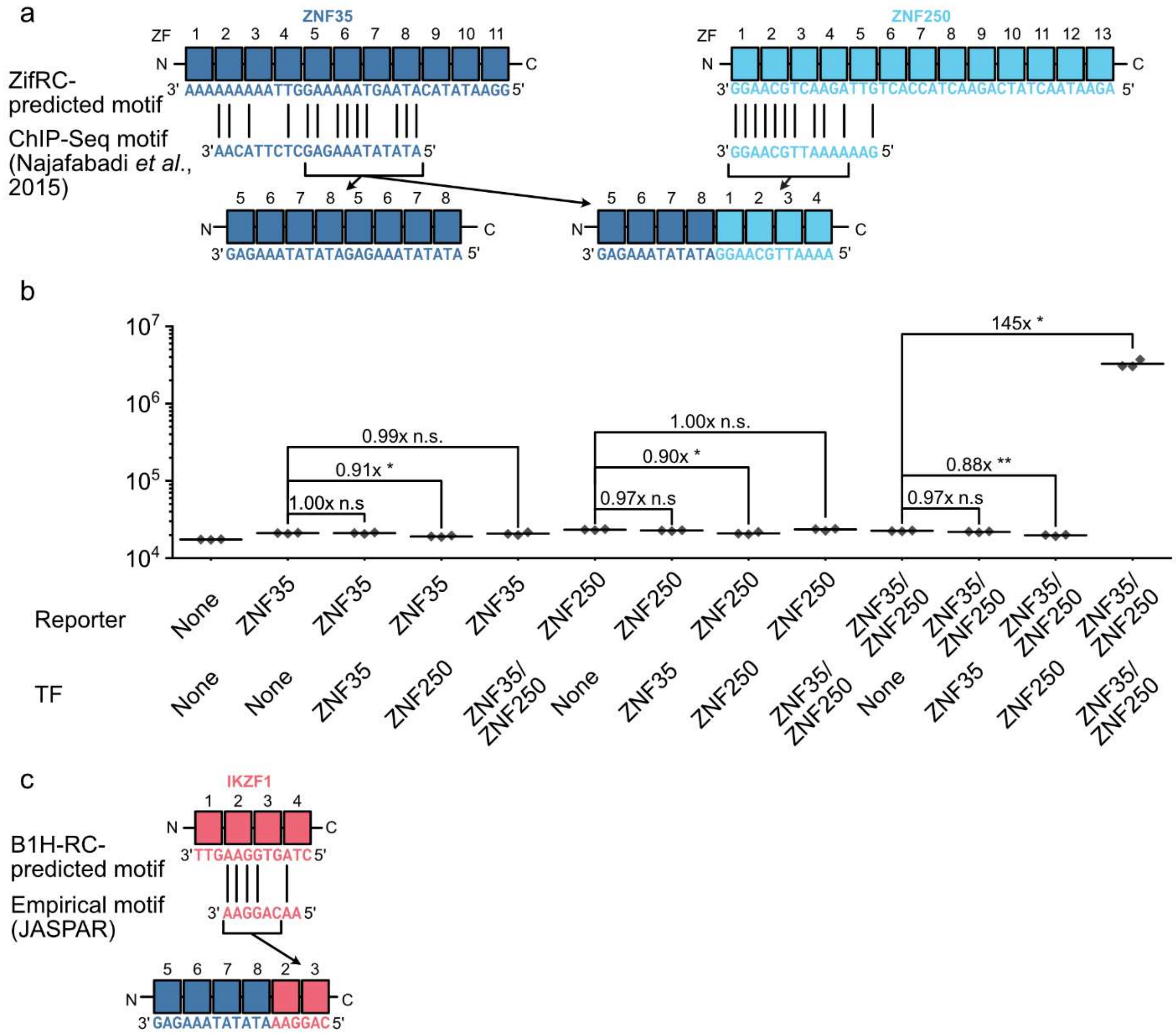


**Supplementary Fig. S1. Mapping of human transcription factor binding motifs to zinc fingers and partial-site binding data for ZNF35/ZNF250 binding domain.** (**a**) Alignment of individual zinc fingers in ZNF35 and ZNF250 to those proteins' empirical binding motifs to generate the binding motifs of the ZNF35/ZNF35 and ZNF35/ZNF250 block fusion ZF arrays. (**b**) Flow cytometry results from transfection of plasmids encoding ZNF35/ZNF250-NZF transcription factor and a reporter containing four copies of its binding site upstream of a GFP reporter, or combinations of the two containing only the portions derived from either ZNF35 or ZNF250. * denotes $p < 0.05$, ** denotes $p < 0.005$, and n.s. denotes $p > 0.05$. (**c**) Alignment of individual zinc fingers in IKZF1 to its empirical binding motifs to generate the binding motif of the ZNF35/IKZF1 block fusion ZF array.

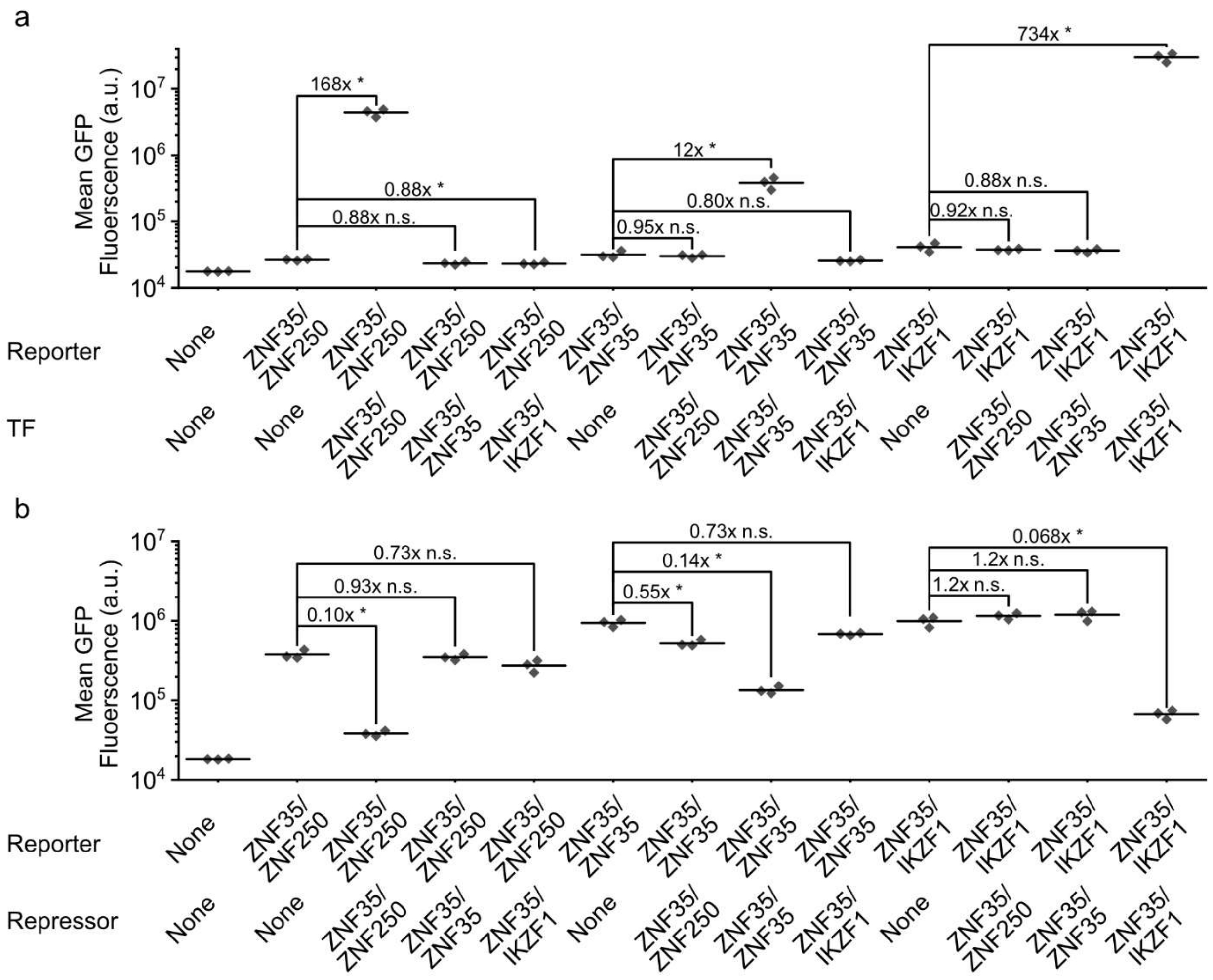


**Supplementary Fig. S2. Complete data for transcription factor and repressor orthogonality matrices.** (**a**) Flow cytometry results from transfection of HEK293 cells with plasmids encoding transcription factors containing each block fusion zinc finger array joined to the NZF transcriptional activation domain along with reporters containing four copies of their corresponding binding sites upstream of a GFP reporter. * denotes $p < 0.05$ and n.s. denotes $p > 0.05$. (**b**) Flow cytometry results from transfection of HEK293 cells with plasmids encoding transcriptional repressors containing each block fusion zinc finger array joined to the ZIK1 KRAB domains domain along with reporters containing four copies of their corresponding binding sites upstream of an SFFV promoter-driven GFP reporter. * denotes $p < 0.05$ and n.s. denotes $p > 0.05$.

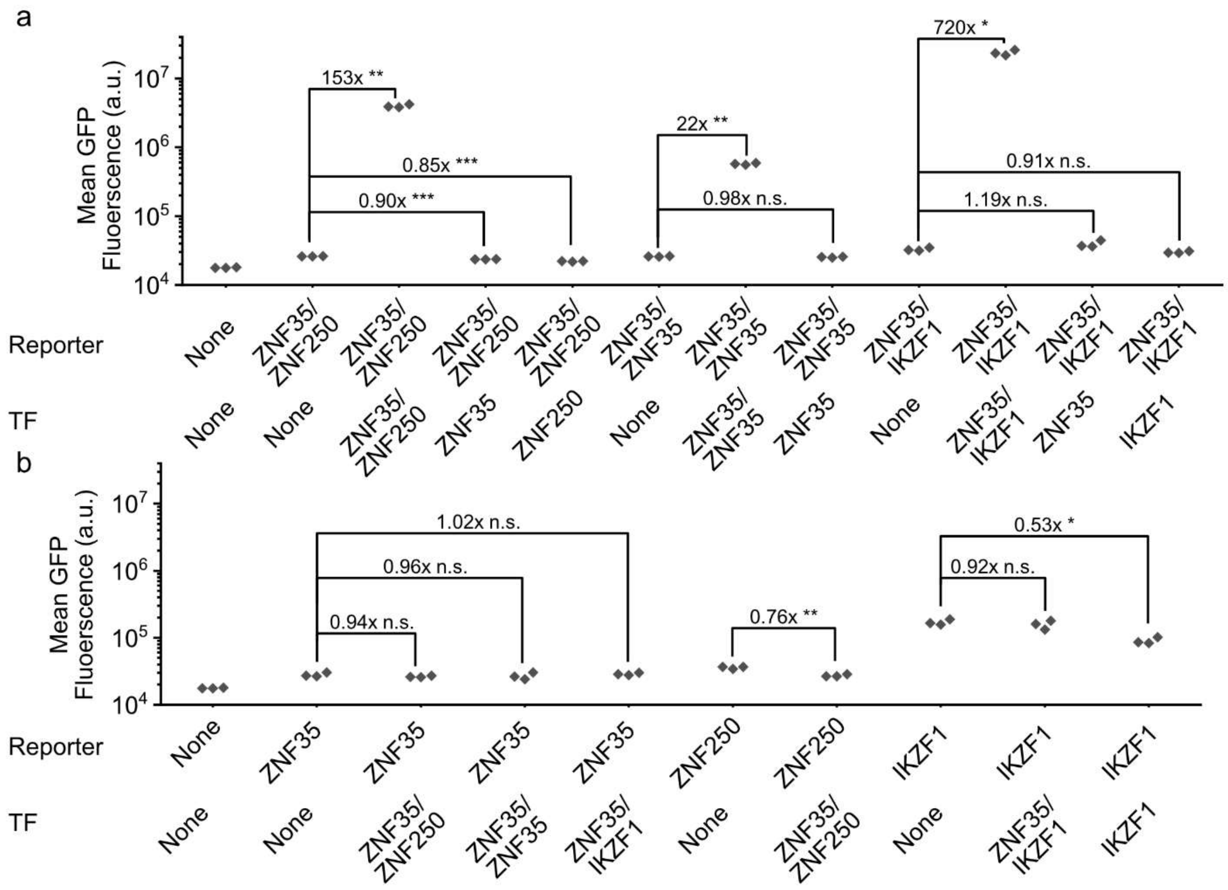


**Supplementary Fig. S3. Measurement of partial binding domain and partial reporter target sequence activity.** (**a**) Flow cytometry results from transfection of HEK293 cells with plasmids encoding either transcription factors containing each block fusion zinc finger array or joined to the NZF transcriptional activation domain or versions containing the partial binding domain from a single protein along with reporters containing four copies of their corresponding full binding sites upstream of a GFP reporter. * denotes $p < 0.05$, ** denotes $p < 0.005$, *** denotes $p < 0.0005$, and n.s. denotes $p > 0.05$. (**b**) Flow cytometry results from transfection of HEK293 cells with plasmids encoding transcription factors containing each block fusion zinc finger array joined to the NZF transcriptional activation domain along with reporters containing either four copies of their full binding sites or eight copies of their partial binding sites upstream of a GFP reporter. * denotes $p < 0.05$, ** denotes $p < 0.005$ and n.s. denotes $p > 0.05$.

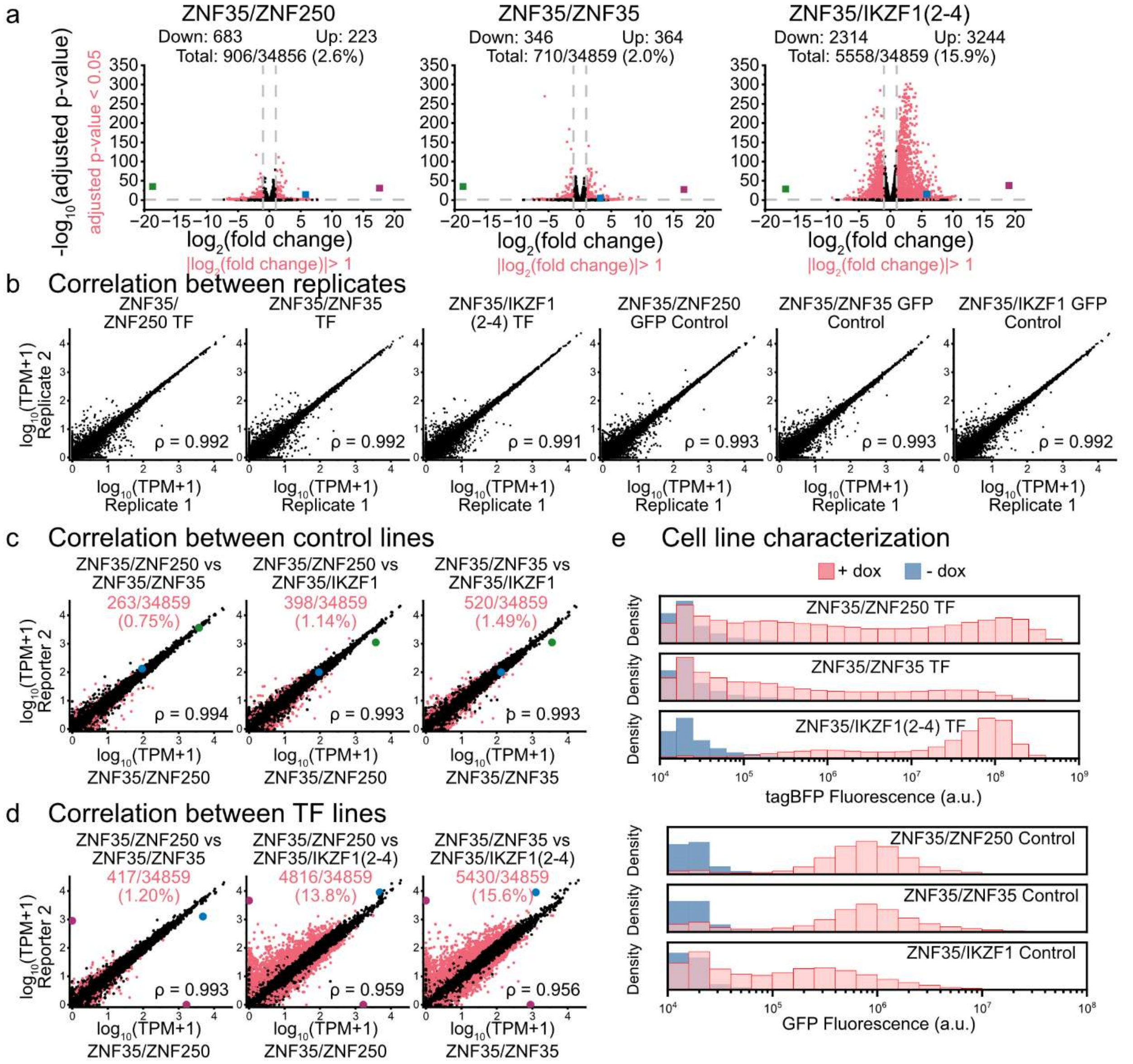


**Supplementary Fig. S4. Complete transcriptomic analysis of stably-integrated transcription factor and control cell lines.** (**a**) Differential expression analysis of cell lines with integrated block fusion ZF transcription factors compared to GFP controls. Red points indicate transcripts with significantly different expression (|$\log_2$(Fold Change| > 2 and adjusted p-value < 0.05). Grey lines indicate thresholds for determination of differential expression. Large green, blue, and purple points indicate transcripts for GFP-NZF, H2B-tagBFP, and the indicated synthetic transcription factor respectively. Numbers of upregulated, downregulated, and total differentially expressed genes are indicated above each graph. (**b**) Correlation of measured TPM values between technical replicates of each cell line. (**c**) Correlation of measured TPM values between GFP-NZF control cell lines. (**d**) Correlation of measured TPM values between cell lines with different block fusion transcription factors integrated. (**e**) Flow cytometric characterization of cell lines by tagBFP fluorescence for transcription factor-integrated cell lines and GFP fluorescence for GFP control cell lines. Cells in “+ dox” conditions were incubated with media containing 100 ng/mL doxycycline for 48 hours prior to flow cytometry. Cells were gated for scattering to exclude noncellular debris but no other gating was performed.

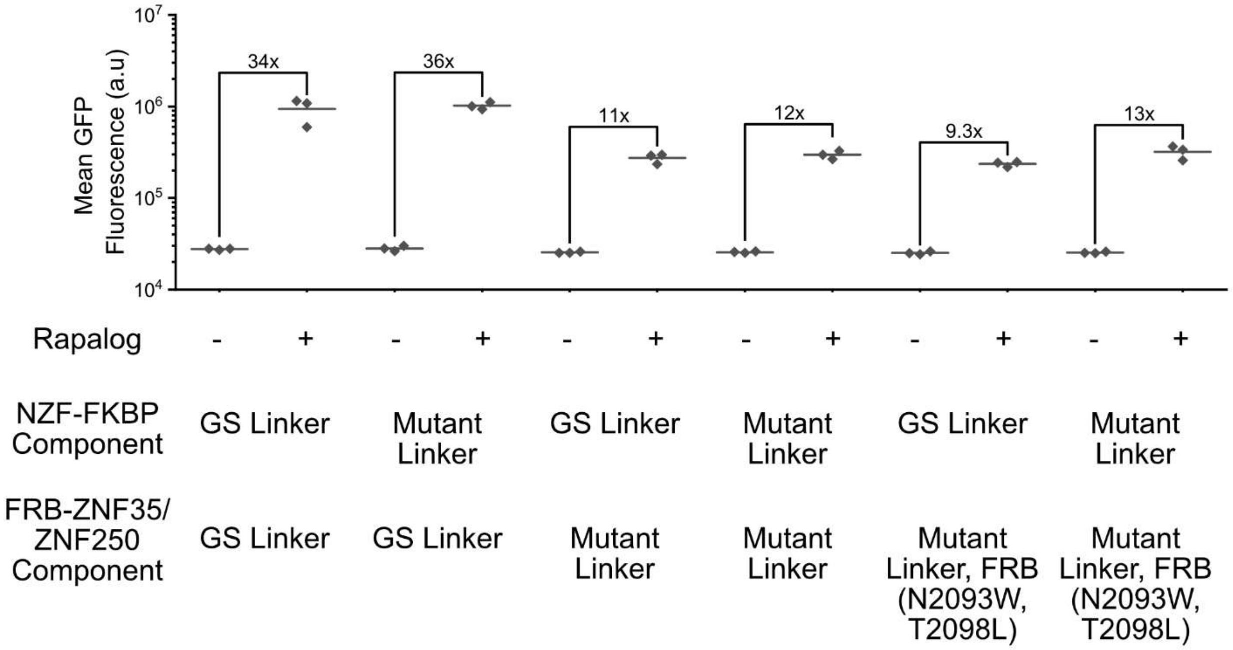


**Supplementary Fig. S5. Rapalog sensitivity of split transcription factors with progressively deimmunized linkers and deimmunized FRB domain.** All conditions are co-transfected with a reporter containing four copies of the ZNF35/ZNF250 binding site upstream of GFP. “+ Rap” conditions received 500 nM rapalog; “- Rap” conditions received 1/1000 ethanol vehicle. Amino acid sequences of all constructs are found in **Supp. Table S1**.

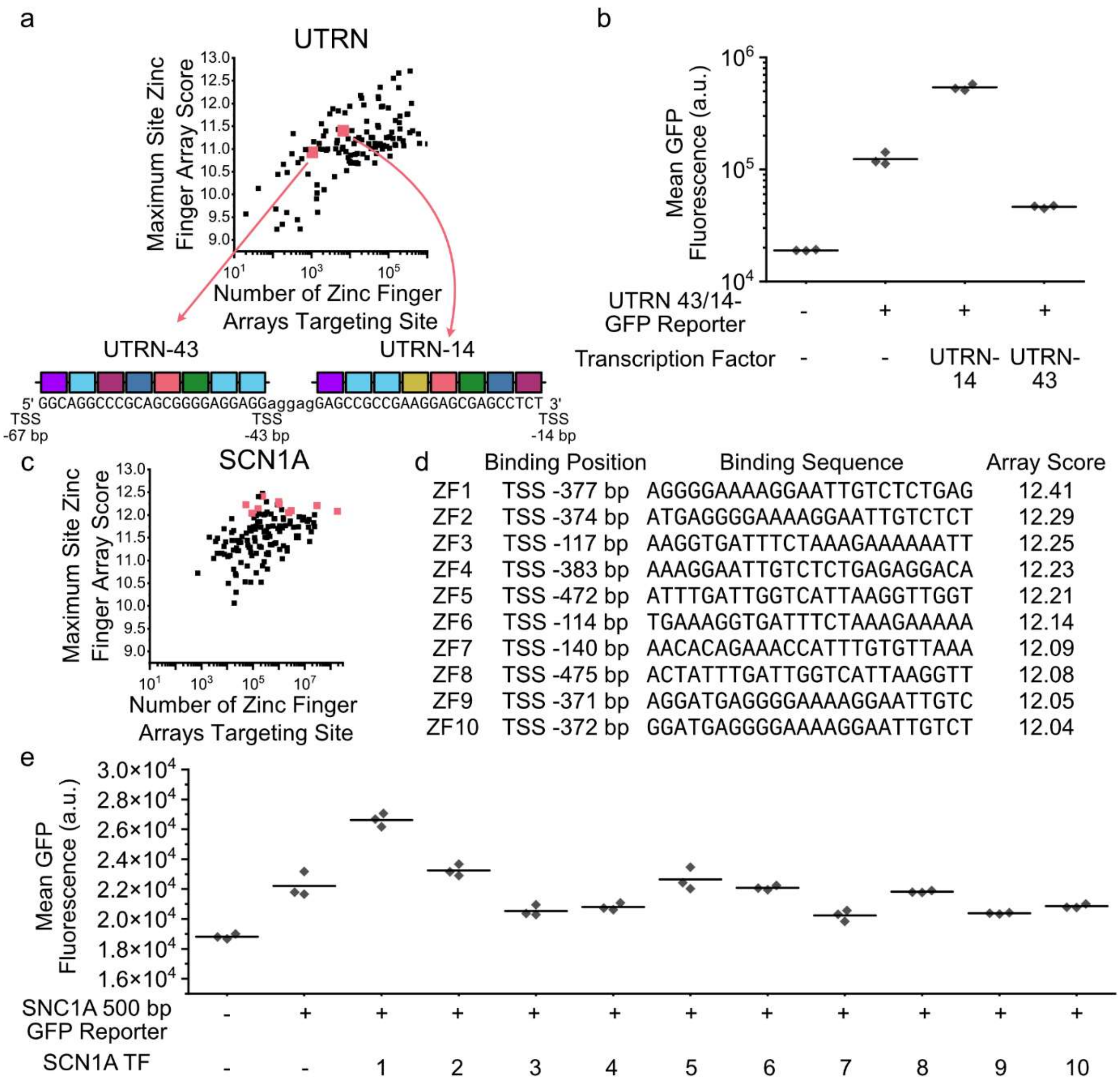


**Supplementary Fig. S6. Screening of UTRN and SCN1A-targeting DNA-binding domains.** (**a**) Identification of UTRN-14 and UTRN-43 arrays and corresponding binding sites. (**b**) Flow cytometry results of co-transfecting a GFP reporter containing one copy of the joint UTRN-43/UTRN-14 binding site with UTRN-14 or UTRN-43 attached to the VP64 transcriptional activation domain. (**c**) Identification of screened SCN1A promoter sites. Red sites denote sites with ZF arrays selected for screening. (**d**) Description of screened UTRN-targeting ZFs and their corresponding binding sites. (**e**) Flow cytometry results of co-transfecting a GFP reporter containing the 500 bp region upstream of the SCN1A transcriptional start with each of the 10 screened SCN1A-targeting ZFs attached to the NZF transcriptional activation domain.

**Supplementary Table S1. Fusion Protein Sequences**

| Protein | Sequence | Key |
|---|---|---|
| Gal4-NZF (GS Linkers) | KLLSSIEQACDICRLKKLKCSKEKPKCAKCL<br>KNNWECRYSPKTKRSPLTRAHLTEVESRLE<br>RLEQLFLLIFPREDLDMILKMDSLQDIKALLT<br>GLFVQDNVNKDAVTDRLASVETDMPLTLRQ<br>HRISATSSSEESSNKGQRQLTVSAAAGGSG<br>GSGGSEGQSDERALLDQLHTLLSNTDATGL<br>EEIDRALGIPELVNQGQALEPKQDGGSGSM<br>AEEFVTLKDVGMDFTLGDWEQLGLEQGDT<br>FWDTALDNCQDLFLLDPPGGGSGGGSGHE<br>KFPSDLDLDMFNGSLECDMESIIRSELMDA<br>DGLDFNFDS | Gal4<br>NCOA3 TAD<br>ZNF473 KRAB<br>FOXO3 TAD |
| Gal4-NZF (Mutated Linkers) | KLLSSIEQACDICRLKKLKCSKEKPKCAKCL<br>KNNWECRYSPKTKRSPLTRAHLTEVESRLE<br>RLEQLFLLIFPREDLDMILKMDSLQDIKALLT<br>GLFVQDNVNKDAVTDRLASVETDMPLTLRQ<br>HRISATSSSEESSNKGQRQLTVSAAAGGSG<br>GSGGSEGQSDERALLDQLHTLLSNTDATGL<br>EEIDRALGIPELVNQGQALEPKQDGGWGS<br>MAEEFVTLKDVGMDFTLGDWEQLGLEQGD<br>TFWDTALDNCQDLFLLDPPGGGDGLGLGH<br>EKFPSDLDLDMFNGSLECDMESIIRSELMD<br>ADGLDFNFDS | Gal4<br>NCOA3 TAD<br>ZNF473 KRAB<br>FOXO3 TAD<br>Mutated residue |
| ADAR2dd (E488Q, T501A)-PUF-9R | LHLDQTPSRQPIPSEGLQLHLPQVLADAVS<br>RLVLGKFGDLTDNFSSPHARRKVLAGVVMT<br>TGTDVKDAKVISVSTGTKCINGEYMSDRGL<br>ALNDCHAEIISRRSLLRFLYTQLELYLNNKDD<br>QKRSIFQKSERGGFRLKENVQFHLYISTSPC<br>GDARIFSPHEPILEEPADRHPNRKARGQLR<br>TKIESGQGTIPVRSNASIQAWDGVLQGERL<br>LTMSCSDKIARWNVVGIQGSLLSIFVEPIYFS<br>SIILGSLYHGDHLSRAMYQRISNIEDLPPLYT<br>LNKPLLSGISNAEARQPGKAPNFSVNWTVG<br>DSAIEVINATTGKDELGRASRLCKHALYCRW<br>MRVHGKVPSHLLRSKITKPNVYHESKLAAK<br>EYQAAKARLFTAFIKAGLGAWVEKPTEQDQ<br>FSLTPGTGGGTGGGAGRSRLLEDFRNNRY<br>PNLQLREIAGHIMEFSQDQHGSRFIQLKLER<br>ATPAERQLVFNEILQAAYQLMVDVFGNYVIQ<br>KFFEFGSLEQKLALAERIRGHVLSLALQMY<br>GCRVIQKALEFIPSDQQNEMVRELHQHTEQ<br>LVQDQYGNYVIQHVLEHGRPEDKSKIVAEIR<br>GNVLVLSQHKFASNVVEKCVTHASRTERAV<br>LIDEVCTMNDGPHSALYTMMKDQYANYVV<br>QKMIDVAEPGQRKIVMHELHQHTEQLVQDQ<br>YGNYVIQHVLEHGRPEDKSKIVAEIRGNVLV<br>LSQHKFASNVVEKCVTHASRTERAVLIDEV<br>CTMNDGPHSALYTMMKDQYANYVVQKMID | ADAR2dd(E488Q, T501A)<br>PUM1 Repeat 1<br>PUM1 Repeat 2<br>PUM1 Repeat 3<br>PUM1 Repeat 6<br>PUM1 Repeat 7<br>PUM1 Repeat 8 |

| | | |
|---|---|---|
| | VAEPGQRKIVMHKIRPHIATLRKYTYGKHILA<br>KLEKYYMKNGVDLG | |
| ADAR2dd (E488Q, T501A)-PUF-9R (V241D, H243M) | LHLDQTPSRQPIPSEGLQLHLPQVLADAVS<br>RLVLGKFGDLTDNFSSPHARRKVLAGVVMT<br>TGTDVKDAKVISVSTGTKCINGEYMSDRGL<br>ALNDCHAEIISRRSLLRFLYTQLELYLNNKDD<br>QKRSIFQKSERGGFRLKENVQFHLYISTSPC<br>GDARIFSPHEPILEEPADRHPNRKARGQLR<br>TKIESGQGTIPVRSNASIQAWDGVLQGERL<br>LTMSCSDKIARWNVVGIQGSLLSIFVEPIYFS<br>SIILGSLYHGDHLSRAMYQRISNIEDLPPLYT<br>LNKPLLSGISNAEARQPGKAPNFSVNWTVG<br>DSAIEVINATTGKDELGRASRLCKHALYCRW<br>MRVHGKVPSHLLRSKITKPNVYHESKLAAK<br>EYQAAKARLFTAFIKAGLGAWVEKPTEQDQ<br>FSLTPGTGGGTGGGAGRSRLLEDFRNNRY<br>PNLQLREIAGHIMEFSQDQHGSRFIQLKLER<br>ATPAERQLVFNEILQAAYQLMVDVFGNYVIQ<br>KFFEFGSLEQKLALAERIRGHVLSLALQMY<br>GCRVIQKALEFIPSDQQNEMVRELHQHTEQ<br>LVQDQYGNYVIQHVLEHGRPEDKSKIVAEIR<br>GNVLVLSQHKFASNVVEKCVTHASRTERAV<br>LIDEVCTMNDGPHSALYTMMKDQYANYVV<br>QKMIDVAEPGQRKIDMMELHQHTEQLVQD<br>QYGNYVIQHVLEHGRPEDKSKIVAEIRGNVL<br>VLSQHKFASNVVEKCVTHASRTERAVLIDEV<br>CTMNDGPHSALYTMMKDQYANYVVQKMID<br>VAEPGQRKIVMHKIRPHIATLRKYTYGKHILA<br>KLEKYYMKNGVDLG | ADAR2dd(E488Q, T501A)<br>PUM1 Repeat 1<br>PUM1 Repeat 2<br>PUM1 Repeat 3<br>PUM1 Repeat 6<br>PUM1 Repeat 7<br>PUM1 Repeat 8<br>Mutated residue |
| ADAR2dd (E488Q, T501A)-PUF-9R(K239S,V241L, H243M) | LHLDQTPSRQPIPSEGLQLHLPQVLADAVS<br>RLVLGKFGDLTDNFSSPHARRKVLAGVVMT<br>TGTDVKDAKVISVSTGTKCINGEYMSDRGL<br>ALNDCHAEIISRRSLLRFLYTQLELYLNNKDD<br>QKRSIFQKSERGGFRLKENVQFHLYISTSPC<br>GDARIFSPHEPILEEPADRHPNRKARGQLR<br>TKIESGQGTIPVRSNASIQAWDGVLQGERL<br>LTMSCSDKIARWNVVGIQGSLLSIFVEPIYFS<br>SIILGSLYHGDHLSRAMYQRISNIEDLPPLYT<br>LNKPLLSGISNAEARQPGKAPNFSVNWTVG<br>DSAIEVINATTGKDELGRASRLCKHALYCRW<br>MRVHGKVPSHLLRSKITKPNVYHESKLAAK<br>EYQAAKARLFTAFIKAGLGAWVEKPTEQDQ<br>FSLTPGTGGGTGGGAGRSRLLEDFRNNRY<br>PNLQLREIAGHIMEFSQDQHGSRFIQLKLER<br>ATPAERQLVFNEILQAAYQLMVDVFGNYVIQ<br>KFFEFGSLEQKLALAERIRGHVLSLALQMY<br>GCRVIQKALEFIPSDQQNEMVRELHQHTEQ<br>LVQDQYGNYVIQHVLEHGRPEDKSKIVAEIR<br>GNVLVLSQHKFASNVVEKCVTHASRTERAV<br>LIDEVCTMNDGPHSALYTMMKDQYANYVV<br>QKMIDVAEPGQRSILMMELHQHTEQLVQDQ | ADAR2dd(E488Q, T501A)<br>PUM1 Repeat 1<br>PUM1 Repeat 2<br>PUM1 Repeat 3<br>PUM1 Repeat 6<br>PUM1 Repeat 7<br>PUM1 Repeat 8<br>Mutated residue |

| | | |
|---|---|---|
| | YGNYVIQHVLEHGRPEDKSKIVAEIRGNVLV<br>LSQHKFASNVVEKCVTHASRTERAVLIDEV<br>CTMNDGPHSALYTMMKDQYANYVVQKMID<br>VAEPGQRKIVMHKIRPHIATLRKYTYGKHILA<br>KLEKYYMKNGVDLG | |
| ZNF35(5-8)-<br>ZNF250(1-4)-<br>deImmunLink-NZF | YECNECGKTFTRSSNLIVHQRIHTGEKPFAC<br>NDCGKAFTQSANLIVHQRSHTGEKPYECKE<br>CGKAFSCFSHLIVHQRIHTAEKPYDCSECG<br>KAFSQLSCLIVHQRIHTGERPYMCVECGKC<br>FGRSSHLLQHQRIHTGEKPYVCSVCGKAFS<br>QSSVLSKHRRIHTGEKPYECNECGKAFRVS<br>SDLAQHHKIHTGEKPHECLECRKAFTQLSH<br>LIQHQRIHAAAWELGMSGGSEGQSDERALL<br>DQLHTLLSNTDATGLEEIDRALGIPELVNQG<br>QALEPKQDGGWGSMAEEFVTLKDVGMDF<br>TLGDWEQLGLEQGDTFWDTALDNCQDLFL<br>LDPPGGGDGLGLGHEKFPSDLDLDMFNGS<br>LECDMESIIRSELMDADGLDFNFDS | ZNF35 (5-8)<br>ZNF250(1-4)<br>NZF (Mutated<br>Linkers) |
| ZNF35(5-8)-<br>ZNF35(5-8)-<br>deImmunLink-NZF | YECNECGKTFTRSSNLIVHQRIHTGEKPFAC<br>NDCGKAFTQSANLIVHQRSHTGEKPYECKE<br>CGKAFSCFSHLIVHQRIHTAEKPYDCSECG<br>KAFSQLSCLIVHQRIHTGERPYECNECGKT<br>FTRSSNLIVHQRIHTGEKPFACNDCGKAFT<br>QSANLIVHQRSHTGEKPYECKECGKAFSCF<br>SHLIVHQRIHTAEKPYDCSECGKAFSQLSCL<br>IVHQRIHAAAWELGMSGGSEGQSDERALL<br>DQLHTLLSNTDATGLEEIDRALGIPELVNQG<br>QALEPKQDGGWGSMAEEFVTLKDVGMDF<br>TLGDWEQLGLEQGDTFWDTALDNCQDLFL<br>LDPPGGGDGLGLGHEKFPSDLDLDMFNGS<br>LECDMESIIRSELMDADGLDFNFDS | ZNF35 (5-8)<br>NZF (Mutated<br>Linkers) |
| ZNF35(5-8)-<br>IKZF1(2-3)-<br>deimmunLink-NZF | YECNECGKTFTRSSNLIVHQRIHTGEKPFAC<br>NDCGKAFTQSANLIVHQRSHTGEKPYECKE<br>CGKAFSCFSHLIVHQRIHTAEKPYDCSECG<br>KAFSQLSCLIVHQRIHTGERPFQCNQCGAS<br>FTQKGNLLRHIKLHSGEKPFKCHLCNYACR<br>RRDALTGHLRTHGWSGGDGMSEGQSDER<br>ALLDQLHTLLSNTDATGLEEIDRALGIPELVN<br>QGQALEPKQDGGWGSMAEEFVTLKDVGM<br>DFTLGDWEQLGLEQGDTFWDTALDNCQDL<br>FLLDPPGGGDGLGLGHEKFPSDLDLDMFN<br>GSLECDMESIIRSELMDADGLDFNFDS | ZNF35 (5-8)<br>IKZF1 (2-3)<br>NZF (Mutated<br>Linkers) |
| ZNF35(5-8)<br>deImmunLink-NZF | YECNECGKTFTRSSNLIVHQRIHTGEKPFAC<br>NDCGKAFTQSANLIVHQRSHTGEKPYECKE<br>CGKAFSCFSHLIVHQRIHTAEKPYDCSECG<br>KAFSQLSCLIVHQRIHAAAWELGMSGGSEG<br>QSDERALLDQLHTLLSNTDATGLEEIDRALG<br>IPELVNQGQALEPKQDGGWGSMAEEFVTL<br>KDVGMDFTLGDWEQLGLEQGDTFWDTALD<br>NCQDLFLLDPPGGGDGLGLGHEKFPSDLD | ZNF35 (5-8)<br>NZF (Mutated<br>Linkers) |

| | | |
|---|---|---|
| | LDMFNGSLECDMESIIRSELMDADGLDFNFDS | |
| ZNF250(1-4)-deImmunLink-NZF | YMCVECGKCFGRSSHLLQHQRIHTGEKPYVCSVCGKAFSQSSVLSKHRRIHTGEKPYECNECGKAFRVSSDLAQHHKIHTGEKPHECLECRKAFTQLSHLIQHQRIHAAAWELGMSGGSEGQSDERALLDQLHTLLSNTDATGLEEIDRALGIPELVNQGQALEPKQDGGWGSMAEEFVTLKDVGMDFTLGDWEQLGLEQGDTFWDTALDNCQDLFLLDPPGGGDGLGLGHEKFPSDLDLDMFNGSLECDMESIIRSELMDADGLDFNFDS | ZNF250(1-4)<br>NZF (Mutated Linkers) |
| IKZF1(2-3)-deimmunLink-NZF | FQCNQCGASFTQKGNLLRHIKLHSGEKPFKCHLCNYACRRRDALTGHLRTHGWSGGDGMSEGQSDERALLDQLHTLLSNTDATGLEEIDRALGIPELVNQGQALEPKQDGGWGSMAEEFVTLKDVGMDFTLGDWEQLGLEQGDTFWDTALDNCQDLFLLDPPGGGDGLGLGHEKFPSDLDLDMFNGSLECDMESIIRSELMDADGLDFNFDS | IKZF1 (2-3)<br>NZF (Mutated Linkers) |
| ZNF35(5-8)-IKZF1(2-4)-deImmunLink-NZF | YECNECGKTFTRSSNLIVHQRIHTGEKPFACNDCGKAFTQSANLIVHQRSHTGEKPYECKECGKAFSCFSHLIVHQRIHTAEKPYDCSECGKAFSQLSCLIVHQRIHTGERPFQCNQCGASFTQKGNLLRHIKLHSGEKPFKCHLCNYACRRRDALTGHLRTHSVGKPHKCGYCGRSYKQRSSLEEHKERCHAAAGWSGGDGGDEGQSDERALLDQLHTLLSNTDATGLEEIDRALGIPELVNQGQALEPKQDGGWGSMAEEFVTLKDVGMDFTLGDWEQLGLEQGDTFWDTALDNCQDLFLLDPPGGGDGLGLGHEKFPSDLDLDMFNGSLECDMESIIRSELMDADGLDFNFDS | ZNF35 (5-8)<br>IKZF1 (2-4)<br>NZF (Mutated Linkers) |
| ZNF35(5-8)-ZNF250(1-4)-deImmunLink -ZIK1KRAB | YECNECGKTFTRSSNLIVHQRIHTGEKPFACNDCGKAFTQSANLIVHQRSHTGEKPYECKECGKAFSCFSHLIVHQRIHTAEKPYDCSECGKAFSQLSCLIVHQRIHTGERPYMCVECGKCFGRSSHLLQHQRIHTGEKPYVCSVCGKAFSQSSVLSKHRRIHTGEKPYECNECGKAFRVSSDLAQHHKIHTGEKPHECLECRKAFTQLSHLIQHQRIHAAAWELGMSGGSCVTFEDIAIYFSQDEWGLLDEAQRLLYLEVMLENFALVASL | ZNF35 (5-8)<br>ZNF250(1-4)<br>ZIK1 KRAB |
| ZNF35(5-8)-ZNF35(5-8)-deImmunLink -ZIK1KRAB | YECNECGKTFTRSSNLIVHQRIHTGEKPFACNDCGKAFTQSANLIVHQRSHTGEKPYECKECGKAFSCFSHLIVHQRIHTAEKPYDCSECGKAFSQLSCLIVHQRIHTGERPYECNECGKTFTRSSNLIVHQRIHTGEKPFACNDCGKAFTQSANLIVHQRSHTGEKPYECKECGKAFSCFSHLIVHQRIHTAEKPYDCSECGKAFSQLSCL | ZNF35 (5-8)<br>ZIK1 KRAB |

| | | |
|---|---|---|
| | IVHQRIHAAAWELGMSGGSCVTFEDIAIYFS<br>QDEWGLLDEAQRLLYLEVMLENFALVASL | |
| ZNF35(5-8)-<br>IKZF1(2-3)-<br>deImmunLink -<br>ZIK1KRAB | YECNECGKTFTRSSNLIVHQRIHTGEKPFAC<br>NDCGKAFTQSANLIVHQRSHTGEKPYECKE<br>CGKAFSCFSHLIVHQRIHTAEKPYDCSECG<br>KAFSQLSCLIVHQRIHTGERPFQCNQCGAS<br>FTQKGNLLRHIKLHSGEKPFKCHLCNYACR<br>RRDALTGHLRTHGWSGGDGMSCVTFEDIAI<br>YFSQDEWGLLDEAQRLLYLEVMLENFALVA<br>SL | ZNF35 (5-8)<br>IKZF1 (2-3)<br>ZIK1 KRAB |
| NZF-FKBP | EGQSDERALLDQLHTLLSNTDATGLEEIDRA<br>LGIPELVNQGQALEPKQDGGWGSMAEEFV<br>TLKDVGMDFTLGDWEQLGLEQGDTFWDTA<br>LDNCQDLFLLDPPGGGDGLGLGHEKFPSD<br>LDLDMFNGSLECDMESIIRSELMDADGLDF<br>NFDSAAAGGSGGSGGSGGSGGSGGSGGS<br>GGSGVQVETISPGDGRTFPKRGQTCVVHY<br>TGMLEDGKKFDSSRDRNKPFKFMLGKQEVI<br>RGWEEGVAQMSVGQRAKLTISPDYAYGAT<br>GHPGIIPPHATLVFDVELLKLE | NZF (Mutated<br>Linkers)<br>FKBP |
| NZF-deImmunLink -<br>FKBP | EGQSDERALLDQLHTLLSNTDATGLEEIDRA<br>LGIPELVNQGQALEPKQDGGWGSMAEEFV<br>TLKDVGMDFTLGDWEQLGLEQGDTFWDTA<br>LDNCQDLFLLDPPGGGDGLGLGHEKFPSD<br>LDLDMFNGSLECDMESIIRSELMDADGLDF<br>NFDSGGMGGMLTTGGTKGMLVSGYSGMS<br>GVQVETISPGDGRTFPKRGQTCVVHYTGM<br>LEDGKKFDSSRDRNKPFKFMLGKQEVIRG<br>WEEGVAQMSVGQRAKLTISPDYAYGATGH<br>PGIIPPHATLVFDVELLKLE | NZF (Mutated<br>Linkers)<br>FKBP<br>Mutated residues |
| FRB(T2098L)-<br>ZNF35(5-8)-<br>ZNF250(1-4) | EMWHEGLEEASRLYFGERNVKGMFEVLEP<br>LHAMMERGPQTLKETSFNQAYGRDLMEAQ<br>EWCRKYMKSGNVKDLLQAWDLYYHVFRRI<br>SKTRAAAGGSGGSGGSYECNECGKTFTRS<br>SNLIVHQRIHTGEKPFACNDCGKAFTQSAN<br>LIVHQRSHTGEKPYECKECGKAFSCFSHLIV<br>HQRIHTAEKPYDCSECGKAFSQLSCLIVHQ<br>RIHTGERPYMCVECGKCFGRSSHLLQHQRI<br>HTGEKPYVCSVCGKAFSQSSVLSKHRRIHT<br>GEKPYECNECGKAFRVSSDLAQHHKIHTGE<br>KPHECLECRKAFTQLSHLIQHQRIH | FRB<br>ZNF35 (5-8)<br>ZNF250(1-4) |
| FRB(T2098L)-<br>deImmunLink-<br>ZNF35(5-8)-<br>ZNF250(1-4) | EMWHEGLEEASRLYFGERNVKGMFEVLEP<br>LHAMMERGPQTLKETSFNQAYGRDLMEAQ<br>EWCRKYMKSGNVKDLLQAWDLYYHVFRRI<br>SKTRGGSGWWVNSYECNECGKTFTRSSN<br>LIVHQRIHTGEKPFACNDCGKAFTQSANLIV<br>HQRSHTGEKPYECKECGKAFSCFSHLIVHQ<br>RIHTAEKPYDCSECGKAFSQLSCLIVHQRIH<br>TGERPYMCVECGKCFGRSSHLLQHQRIHT<br>GEKPYVCSVCGKAFSQSSVLSKHRRIHTGE | FRB<br>ZNF35 (5-8)<br>ZNF250(1-4)<br>Mutated residues |

| | | |
|---|---|---|
| | KPYECNECGKAFRVSSDLAQHHKIHTGEKP HECLECRKAFTQLSHLIQHQRIH | |
| FRB(N2093W, T2098L) deImmunLink-ZNF35(5-8)-ZNF250(1-4) | EMWHEGLEEASRLYFGERNVKGMFEVLEP LHAMMERGPQTLKETSFNQAYGRDLMEAQ EWCRKYMKSGWVKDLLQAWDLYYHVFRRI SKTRGGSGWWVNSYECNECGKTFTRSSN LIVHQRIHTGEKPFACNDCGKAFTQSANLIV HQRSHTGEKPYECKECGKAFSCFSHLIVHQ RIHTAEKPYDCSECGKAFSQLSCLIVHQRIH TGERPYMCVECGKCFGRSSHLLQHQRIHT GEKPYVCSVCGKAFSQSSVLSKHRRIHTGE KPYECNECGKAFRVSSDLAQHHKIHTGEKP HECLECRKAFTQLSHLIQHQRIH | FRB ZNF35 (5-8) ZNF250(1-4) Mutated residues |
| UTRN-14ZF-deImmLink-NZF | HECARCGKNFSWHSDLILHEQIHTGERPFK CYKCGRGFAEHGTLNRHLRTKGGCTGERP YNCEECGRAFSQASHLQDHQRLHTGERPH QCIECGKSFNRHCNLIRHQKIHTGERPFEC SKCGKACTRRCNLIQHQKVHTGERPFACRF CAKPFRRSSDMRDHERVHTGERPFACRFC AKPFRRSSDMRDHERVHTGERPFQCELCS YTCPRRSNLDRHMKSHAGWMGMSGGSEG QSDERALLDQLHTLLSNTDATGLEEIDRALG IPELVNQGQALEPKQDGGWGSMAEEFVTL KDVGMDFTLGDWEQLGLEQGDTFWDTALD NCQDLFLLDPPGGGDGLGLGHEKFPSDLD LDMFNGSLECDMESIIRSELMDADGLDFNF DS | Q5JNZ3 finger 1 Q66K89 finger 10 Q9NYT6 finger 3 O15535 finger 5 Q13398 finger 2 Q8NCA9 finger 4 P49711 finger 1 NZF (Mutated Linkers) |
| UTRN-14ZF (N62K,H89F,K121F) -deImmLink-NZF | HECARCGKNFSWHSDLILHEQIHTGERPFK CYKCGRGFAEHGTLNRHLRTKGGCTGERP YKCEECGRAFSQASHLQDHQRLHTGERPF QCIECGKSFNRHCNLIRHQKIHTGERPFEC SFCGKACTRRCNLIQHQKVHTGERPFACRF CAKPFRRSSDMRDHERVHTGERPFACRFC AKPFRRSSDMRDHERVHTGERPFQCELCS YTCPRRSNLDRHMKSHAGWMGMSGGSEG QSDERALLDQLHTLLSNTDATGLEEIDRALG IPELVNQGQALEPKQDGGWGSMAEEFVTL KDVGMDFTLGDWEQLGLEQGDTFWDTALD NCQDLFLLDPPGGGDGLGLGHEKFPSDLD LDMFNGSLECDMESIIRSELMDADGLDFNF DS | Q5JNZ3 finger 1 Q66K89 finger 10 Q9NYT6 finger 3 O15535 finger 5 Q13398 finger 2 Q8NCA9 finger 4 P49711 finger 1 NZF (Mutated Linkers) Mutated residues |
| SCN1A-377-NZF | YGCKKCGRRFGRLSNCTRHEKTHSACTGE RPHGCHLCGKAFTHCSDLRKHERTHTGER PYSCGICGKSFSDSSAKRRHCILHTGERPF VCGVCKMGFSLLTSLAQHHSSHTGERPYE CKECGKTFRQCSHLKRHQRIHTGERPYQC GQCGKSFRQSSNLHQHHRLHHTGERPYEC KECGKTFRQCSHLKRHQRIHTGERPFECAV CGESFVNPAELADHVTVHAAAGGSGGSGG SEGQSDERALLDQLHTLLSNTDATGLEEIDR | P98182 finger 5 Q8TC21 finger 2 O43167 finger 7 Q9P2F9 finger 2 Q8TAF7 finger 8 Q15697 finger 3 Q9GZU2 finger 9 NZF (Mutated Linkers) |

| | | |
|---|---|---|
| | ALGIPELVNQGQALEPKQDGGWGSMAEEF VTLKDVGMDFTLGDWEQLGLEQGDTFWDT ALDNCQDLFLLDPPGGGDGLGLGHEKFPS DLDLDMFNGSLECDMESIIRSELMDADGLD FNFDS | |
| SCN1A-377(G2E, A104K, S109I)-deImmLink-NZF | YECKKCGRRFGRLSNCTRHEKTHSACTGE RPHGCHLCGKAFTHCSDLRKHERTHTGER PYSCGICGKSFSDSSAKRRHCILHTGERPF VCGVCKMGFSLLTSLKQHHSIHTGERPYEC KECGKTFRQCSHLKRHQRIHTGERPYQCG QCGKSFRQSSNLHQHHRLHHTGERPYECK ECGKTFRQCSHLKRHQRIHTGERPFECAVC GESFVNPAELADHVTVHAAMGMSGGSDGS EGQSDERALLDQLHTLLSNTDATGLEEIDRA LGIPELVNQGQALEPKQDGGWGSMAEEFV TLKDVGMDFTLGDWEQLGLEQGDTFWDTA LDNCQDLFLLDPPGGGDGLGLGHEKFPSD LDLDMFNGSLECDMESIIRSELMDADGLDF NFDS | P98182 finger 5<br>Q8TC21 finger 2<br>O43167 finger 7<br>Q9P2F9 finger 2<br>Q8TAF7 finger 8<br>Q15697 finger 3<br>Q9GZU2 finger 9<br>NZF (Mutated Linkers)<br>Mutated residues |
| UtroUpZF-deImmunLink-NZF | MILDRPYACPVESCDRRFSRSDNLVRHIRIH TGQKPFQCRICMRNFSRSDHLTTHNRTHT GEKPFACDICGRKFADPGHLVRHNRIHTGE KPFACPVESCDRRFSRSDELTRHIRIHTGQK PFQCRICMRNFSSRDVLRRHNRTHTGEKP FACDICGRKFASRDVLRRHNRIHLRQNDLE AAAWELGMSGGSEGQSDERALLDQLHTLL SNTDATGLEEIDRALGIPELVNQGQALEPKQ DGGWGSMAEEFVTLKDVGMDFTLGDWEQ LGLEQGDTFWDTALDNCQDLFLLDPPGGG DGLGLGHEKFPSDLDLDMFNGSLECDMESI IRSELMDADGLDFNFDS | UtroUp<br>NZF (Mutated Linkers) |
| UTRN-14ZF-VP64 | HECARCGKNFSWHSDLILHEQIHTGERPFK CYKCGRGFAEHGTLNRHLRTKGGCTGERP YNCEECGRAFSQASHLQDHQRLHTGERPH QCIECGKSFNRHCNLIRHQKIHTGERPFEC SKCGKACTRRCNLIQHQKVHTGERPFACRF CAKPFRRSSDMRDHERVHTGERPFACRFC AKPFRRSSDMRDHERVHTGERPFQCELCS YTCPRRSNLDRHMKSHAAAGGSGGSGGS DALDDFDLDMLGSDALDDFDLDMLGSDAL DDFDLDMLGSDALDDFDLDMLGS | Q5JNZ3 finger 1<br>Q66K89 finger 10<br>Q9NYT6 finger 3<br>O15535 finger 5<br>Q13398 finger 2<br>Q8NCA9 finger 4<br>P49711 finger 1<br>VP64 |
| UTRN-43ZF-VP64 | FECAVCGESFVNPAELADHVTVHTGERPFE CAVCGESFVNPAELADHVTVHTGERPYEC QECQKTFSRSSHLLRHQSVHCTGERPFTC DSCGFGFSCEKLLDEHVLTCTGERPYECG QCGRYFIQMADFHRHEKCHTGERPYECED CGLGFVDLTDLTDHQKVHTGERPFECAVCG ESFVNPAELADHVTVHTGERPQECPVCHKII HGAGKLPRHMRTHAAAGGSGGSGGSDAL | Q9GZU2 finger 9<br>Q96NJ6 finger 13<br>Q9Y2K1 finger 1<br>Q6ZNG0 finger 1<br>Q9GZU2 finger 6<br>O15156 finger 1<br>VP64 |

| | | |
|---|---|---|
| | DDFDLDMLGSDALDDFDLDMLGSDALDDF<br>DLDMLGSDALDDFDLDMLGS | |
| SCN1A-374-NZF | HGCHLCGKAFTHCSDLRKHERTHTGERPY<br>SCGICGKSFSDSSAKRRHCILHTGERPFVC<br>GVCKMGFSLLTSLAQHHSSHTGERPYECK<br>ECGKTFRQCSHLKRHQRIHTGERPYQCGQ<br>CGKSFRQSSNLHQHHRLHHTGERPYECKE<br>CGKTFRQCSHLKRHQRIHTGERPFECAVC<br>GESFVNPAELADHVTVHTGERPYQCHNCG<br>KSFISKSQLDIHHRIHAAAGGSGGSGGSEG<br>QSDERALLDQLHTLLSNTDATGLEEIDRALG<br>IPELVNQGQALEPKQDGGWGSMAEEFVTL<br>KDVGMDFTLGDWEQLGLEQGDTFWDTALD<br>NCQDLFLLDPPGGGDGLGLGHEKFPSDLD<br>LDMFNGSLECDMESIIRSELMDADGLDFNF<br>DS | Q8TC21 finger 2<br>O43167 finger 7<br>Q9P2F9 finger 2<br>Q8TAF7 finger 8<br>Q15697 finger 3<br>Q9GZU2 finger 9<br>Q9Y473 finger 6<br>NZF (Mutated<br>Linkers) |
| SCN1A-117-NZF | FVCGVCKMGFSLLTSLAQHHSSHTGERPY<br>QCGQCGKSFRQSSNLHQHHRLHHTGERP<br>FICEVCHKSYTQFSNLCRHKRMHADCTGE<br>RPYQCGQCGKSFRQSSNLHQHHRLHHTG<br>ERPHECARCGKNFSWHSDLILHEQIHTGER<br>PFVCGVCKMGFSLLTSLAQHHSSHTGERP<br>YTCEICNKCFTRSAVLRRHKKMHCTGERPF<br>ECSKCGKACTRRCNLIQHQKVHAAAGGSG<br>GSGGSEGQSDERALLDQLHTLLSNTDATGL<br>EEIDRALGIPELVNQGQALEPKQDGGWGS<br>MAEEFVTLKDVGMDFTLGDWEQLGLEQGD<br>TFWDTALDNCQDLFLLDPPGGGDGLGLGH<br>EKFPSDLDLDMFNGSLECDMESIIRSELMD<br>ADGLDFNFDS | Q9P2F9 finger 2<br>Q15697 finger 3<br>Q03112 finger 6<br>Q5JNZ3 finger 1<br>Q6ZSB9 finger 7<br>Q13398 finger 2<br>NZF (Mutated<br>Linkers) |
| SCN1A-383-NZF | YKCNYCGRSYKQQSTLEEHKERCHTGERP<br>FECAVCGESFVNPAELADHVTVHTGERPQE<br>CVECGKSFSRSCNLLRHLLVHTGERPHGC<br>HLCGKAFTHCSDLRKHERTHTGERPYSCGI<br>CGKSFSDSSAKRRHCILHTGERPFVCGVCK<br>MGFSLLTSLAQHHSSHTGERPYACHLCRKA<br>FTQCSHLRRHEKTHTGERPYKCGECGKGF<br>SQSSNLHIHRCIHAAAGGSGGSGGSEGQS<br>DERALLDQLHTLLSNTDATGLEEIDRALGIP<br>ELVNQGQALEPKQDGGWGSMAEEFVTLKD<br>VGMDFTLGDWEQLGLEQGDTFWDTALDN<br>CQDLFLLDPPGGGDGLGLGHEKFPSDLDL<br>DMFNGSLECDMESIIRSELMDADGLDFNFD<br>S | Q9H2S9 finger 4<br>Q9GZU2 finger 9<br>Q96SZ4 finger 13<br>Q8TC21 finger 2<br>O43167 finger 7<br>Q9P2F9 finger 2<br>A8MUZ8 finger 2<br>Q16600 finger 6<br>NZF (Mutated<br>Linkers) |
| SCN1A-472-NZF | YKCNECGKTFNVQSHLSRHHRLHTGERPY<br>DCRECGKAFSHRSSLSRHLMSHTGERPFE<br>CSKCGKACTRRCNLIQHQKVHTGERPFVC<br>GVCKMGFSLLTSLAQHHSSHTGERPYECQ<br>ECGKAFHSPRSCHRHERSHTGERPHKCAV<br>CGFTTENLLQFHEHIPQHTGERPFQCALCQ | Q96N58 finger 9<br>Q9BSG1 finger 2<br>Q13398 finger 2<br>Q9P2F9 finger 2<br>Q8NDP4 finger 4<br>Q9HCE3 finger 8 |

| | | |
|---|---|---|
| | KSFTQLAHLQKHHLVHTGERPYQCNECGK<br>SFRVHSSLGIHQRIHAAAGGSGGSGGSEG<br>QSDERALLDQLHTLLSNTDATGLEEIDRALG<br>IPELVNQGQALEPKQDGGWGSMAEEFVTL<br>KDVGMDFTLGDWEQLGLEQGDTFWDTALD<br>NCQDLFLLDPPGGGDGLGLGHEKFPSDLD<br>LDMFNGSLECDMESIIRSELMDADGLDFNF<br>DS | Q8IZ20 finger 2<br>P51786 finger 8<br>NZF (Mutated<br>Linkers) |
| SCN1A-114-NZF | YQCGQCGKSFRQSSNLHQHHRLHHTGER<br>PFICEVCHKSYTQFSNLCRHKRMHADCTGE<br>RPYQCGQCGKSFRQSSNLHQHHRLHHTG<br>ERPHECARCGKNFSWHSDLILHEQIHTGER<br>PFVCGVCKMGFSLLTSLAQHHSSHTGERP<br>YTCEICNKCFTRSAVLRRHKKMHCTGERPF<br>ECSKCGKACTRRCNLIQHQKVHTGERPFQ<br>CALCQKSFTQLAHLQKHHLVHAAAGGSGG<br>SGGSEGQSDERALLDQLHTLLSNTDATGLE<br>EIDRALGIPELVNQGQALEPKQDGGWGSM<br>AEEFVTLKDVGMDFTLGDWEQLGLEQGDT<br>FWDTALDNCQDLFLLDPPGGGDGLGLGHE<br>KFPSDLDLDMFNGSLECDMESIIRSELMDA<br>DGLDFNFDS | Q15697 finger 3<br>Q03112 finger 6<br>Q5JNZ3 finger 1<br>Q9P2F9 finger 2<br>Q6ZSB9 finger 7<br>Q13398 finger 2<br>Q8IZ20 finger 2<br>NZF (Mutated<br>Linkers) |
| SCN1A-140-NZF | YKCKECGKAFKQYSNLPQHKRTHTGERPY<br>DCRECGKAFSHRSSLSRHLMSHTGERPYA<br>CGECGEAFAWLSHLMEHHSSHTGERPFVC<br>GVCKMGFSLLTSLAQHHSSHTGERPYECQ<br>ECGERFICGSTLKCHESVHTGERPFICEVC<br>HKSYTQFSNLCRHKRMHADCTGERPYKCN<br>YCGRSYKQQSTLEEHKERCHTGERPWQC<br>RICEDMFDSQEYVKQHCMSLASHAAAGGS<br>GGSGGSEGQSDERALLDQLHTLLSNTDAT<br>GLEEIDRALGIPELVNQGQALEPKQDGGWG<br>SMAEEFVTLKDVGMDFTLGDWEQLGLEQG<br>DTFWDTALDNCQDLFLLDPPGGGDGLGLG<br>HEKFPSDLDLDMFNGSLECDMESIIRSELM<br>DADGLDFNFDS | Q3SXZ3 finger 11<br>Q9BSG1 finger 2<br>Q8TBC5 finger 1<br>Q9P2F9 finger 2<br>Q8WTR7 finger 8<br>Q03112 finger 6<br>Q9H2S9 finger 4<br>Q9Y4E5 finger 7<br>NZF (Mutated<br>Linkers) |
| SCN1A-475-NZF | YECTKCRTVFTHLSSLKRHVKSHCTGERPF<br>ECSKCGKACTRRCNLIQHQKVHTGERPFV<br>CGVCKMGFSLLTSLAQHHSSHTGERPYEC<br>QECGKAFHSPRSCHRHERSHTGERPHKCA<br>VCGFTTENLLQFHEHIPQHTGERPFQCALC<br>QKSFTQLAHLQKHHLVHTGERPFFCGECG<br>KAFSCHSSLNVHQRIHTGERPYHCKECGKS<br>FTVGSTLLQHQQIHAAAGGSGGSGGSEGQ<br>SDERALLDQLHTLLSNTDATGLEEIDRALGI<br>PELVNQGQALEPKQDGGWGSMAEEFVTLK<br>DVGMDFTLGDWEQLGLEQGDTFWDTALD<br>NCQDLFLLDPPGGGDGLGLGHEKFPSDLD<br>LDMFNGSLECDMESIIRSELMDADGLDFNF<br>DS | Q68EA5 finger 1<br>Q13398 finger 2<br>Q9P2F9 finger 2<br>Q8NDP4 finger 4<br>Q9HCE3 finger 8<br>Q8IZ20 finger 2<br>Q16587 finger 3<br>A8MQ14 finger 13<br>NZF (Mutated<br>Linkers) |

| | | |
|---|---|---|
| SCN1A-371-NZF | YSCGICGKSFSDSSAKRRHCILHTGERPFVCGVCKMGFSLLTSLAQHHSSHTGERPYECKECGKTFRQCSHLKRHQRIHTGERPYQCGQCGKSFRQSSNLHQHHRLHHTGERPYECKECGKTFRQCSHLKRHQRIHTGERPFECAVCGESFVNPAELADHVTVHTGERPYQCLECGQLLMSPSQLLEHQELHTGERPFECAVCGESFVNPAELADHVTVHAAAGGSGGSGGSEGQSDERALLDQLHTLLSNTDATGLEEIDRALGIPELVNQGQALEPKQDGGWGSMAEEFVTLKDVGMDFTLGDWEQLGLEQGDTFWDTALDNCQDLFLLDPPGGGDGLGLGHEKFPSDLDLDMFNGSLECDMESIIRSELMDADGLDFNFDS | O43167 finger 7<br>Q9P2F9 finger 2<br>Q8TAF7 finger 8<br>Q15697 finger 3<br>Q9GZU2 finger 9<br>Q6ZN55 finger 2<br>NZF (Mutated Linkers) |
| SCN1A-372-NZF | HGCHLCGKAFTHCSDLRKHERTHTGERPYECKECGKAFICGYQLTLHLRTHTGERPFICEVCHKSYTQFSNLCRHKRMHADCTGERPFECSKCGKACTRRCNLIQHQKVHTGERPFICEVCHKSYTQFSNLCRHKRMHADCTGERPYNCEECGKAFNRCSHLTRHKKIHTGERPFQCALCQKSFTQLAHLQKHHLVHTGERPYTCHLCRKAFTQCSHLRRHEKTHAAAGGSGGSGGSEGQSDERALLDQLHTLLSNTDATGLEEIDRALGIPELVNQGQALEPKQDGGWGSMAEEFVTLKDVGMDFTLGDWEQLGLEQGDTFWDTALDNCQDLFLLDPPGGGDGLGLGHEKFPSDLDLDMFNGSLECDMESIIRSELMDADGLDFNFDS | Q8TC21 finger 2<br>Q86UE3 finger 10<br>Q03112 finger 6<br>Q13398 finger 2<br>O95780 finger 10<br>Q8IZ20 finger 2<br>A8MWA4 finger 1<br>NZF (Mutated Linkers) |
